# Gene-specific exponent-corrected normalization for library size in bulk RNA-seq

**DOI:** 10.64898/2026.07.04.736167

**Authors:** RuoFei Yin, DanYang Li, Wei Zong, Kyle D Ketchesin, Marianne L Seney, Colleen A McClung, Pedro L. Baldoni, George C Tseng

## Abstract

Correcting for library size is an essential step in bulk RNA-seq analyses, as differences in sequencing depth across samples can obscure biological signal with technical noise. While numerous normalization methods and model-based strategies have been proposed, we demonstrate here that library size-normalized counts and differential expression results obtained from such widely adopted approaches often remain strongly correlated with library size in large-scale RNA-seq experiments. Through a systematic analysis of over 100 publicly available GEO and TCGA RNA-seq datasets with raw count data, we show that library size association is observed for a substantial proportion of genes even after state-of-the-art library size correction approaches recommended by leading normalization tools. To address this issue, we propose *gecco*, a gene-specific exponent-corrected normalization method for RNA-seq counts that incorporates library size directly into the statistical framework via a gene-specific correction term, rather than applying a uniform adjustment factor across all genes. This formulation generalizes existing normalization approaches and yields normalized counts that are free of residual library size effects. Using both simulation studies and real large-scale RNA-seq datasets, we show that our method mitigates library size bias while preserving biological signal across a range of parameter settings. We further demonstrate that our approach leads to higher detection accuracy and more biologically meaningful pathway enrichment results in downstream differential expression and rhythmicity analyses without compromising false discovery rate control. Our method is implemented in R and is fully compatible with the widely used differential expression analysis methods *DESeq2* and *edgeR*.

## Introduction

Short-read RNA sequencing (RNA-seq) has revolutionized biomedical research by enabling high-throughput transcriptome profiling without the probe libraries required by microarray-based methods. Compared with earlier technologies, RNA-seq provides greater accuracy, sensitivity, and the ability to detect novel RNA isoforms, offering a comprehensive view of gene expression [1]. RNA-seq has become a fundamental tool across many fields of biomedical research, including cancer biology, immunology, developmental biology, and psychiatry, with widely performed downstream analyses that include differential expression (DE) analysis, rhythmicity analysis, and disease subtyping [2–7].

RNA-seq measurements are affected by both biological and technical variation, which must be distinguished from one another for accurate downstream inference [8]. Biological variability arises from genuine differences in gene expression across conditions, individuals, or cell types. In contrast, technical variability can be introduced at multiple stages of the experimental protocol and computational workflow, such as during library preparation, sequencing and read alignment. Differences in the total number of sequenced reads between samples, commonly referred to as sequencing depth or library size (hereafter library size), represent a major source of technical variability that influences observed count sizes and must be accounted for in DE analyses [9, 10]. Technical variability can also be introduced in downstream analyses when molecular features such as GC content, RNA degradation, fragment size distribution, and ambiguity in assigning reads vary systematically across samples [11–14]. If unaccounted for, these sources of technical variation can introduce bias and reduce statistical power in downstream analyses such as DE analysis.

Numerous strategies have been developed to adjust for library size differences in RNA-seq data analysis. Common transformations such as counts per million (CPM) and reads per kilobase per million reads (RPKM, or FPKM if fragment-level counting is performed) adjust counts for library size and are widely used in exploratory data analyses and by certain linear model-based differential analysis methods [6, 15]. Other count-based DE methods, such as *DESeq2* and *edgeR*, adjust for library size in a model-based manner, accounting for RNA composition effects whereby a small set of highly expressed genes can dominate total read counts. Two widely used approaches are the median ratio method in *DESeq2* [4, 16] and the trimmed mean of M-values (TMM) method in *edgeR* [3, 17]. Additional methods account for technical biases beyond library size, including GC content and gene length. For example, EDASeq [18] applies within-lane and between-lane normalization to correct for GC content biases and library size differences, whereas CQN [19] models GC content and gene length effects through gene- and sample-specific offsets.

In this article, we identify a systematic issue in library size normalization for large-scale RNA-seq experiments. In principle, normalization procedures should remove associations between expression measurements and library size. However, we found that across 105 publicly available GEO and TCGA datasets with raw count data, normalized expression values often remain strongly associated with library size. Similar patterns are also observed in downstream DE analyses, where library size-associated DE signals frequently persist despite model-based normalization procedures. The issue reported in this article is not unique to bulk RNA-seq and analogous observations have been reported in single-cell RNA-seq (scRNA-seq) studies [20], although concerns regarding model overfitting have also been raised [21].

In this manuscript, we present *gecco*, a <u>g</u>ene-specific <u>e</u>xponent-<u>c</u>orrected for RNA-seq <u>co</u>unts normalization strategy to address library size artifacts in bulk RNA-seq analyses. We propose a generalized linear model (GLM) framework in which library size is modeled as a gene-specific estimable parameter rather than a fixed offset, allowing direct integration into popular DE methods such as *DESeq2* and *edgeR*, while also providing normalized counts for downstream analyses such as rhythmicity analysis. Using simulation studies and large-scale public RNA-seq datasets, we show that *gecco* effectively reduces library size artifacts while preserving biological signal. We further demonstrate its advantages in DE and rhythmicity analyses, where *gecco* improves detection accuracy and pathway enrichment results.

## Materials and methods

### Generalized linear model for RNA-seq data

We briefly review the negative binomial generalized linear model (NB-GLM) used by *DESeq2* and *edgeR* to model RNA-seq counts. In RNA-seq experiments, we observe counts *y*_*gi*_ for *G* genes, *g* = 1, …, *G*, and *n* RNA samples, *i* = 1, …, *n*. Let *π*_*gi*_ denote the true expression of gene *g* in sample *i*, which, in the context of RNA-seq read counting, refers to the unobserved fraction of nucleotides originating from gene *g* in sample *i*, such that 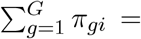 for every sample *i*. For a given gene, it is reasonable to assume that cDNA fragments are subject to Poisson sampling prior to read counting [10]. This assumption implies that, upon sequencing 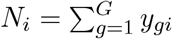 fragments in sample *i*, we have *E*(*Y*_*gi*_|*π*_*gi*_) = *N*_*i*_*π*_*gi*_ and *V ar*(*Y*_*gi*_|*π*_*gi*_) = *N*_*i*_*π*_*gi*_.

The underlying expression level of each gene varies among biological replicates, and a hierarchical model is commonly used to account for this source of *biological* variation. Specifically, we assume that, conditional on the experimental design (which we make implicit to facilitate exposition), the *π*_*gi*_ have a global mean *E*(*π*_*gi*_) = *π*_0,*gi*_ and a constant coefficient of variation for each gene, such that 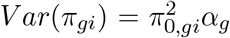 where *α*_*g*_ represents the gene-specific dispersion parameter. Using the law of total variance, it follows that

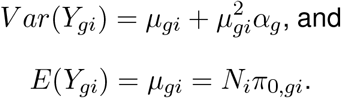

From the above derivation, it follows that any modelling of RNA-seq counts must account for the library size *N*_*i*_, or a proxy thereof, in the mean model.

Both *edgeR* and *DESeq2* model the mean count using a log-linear model with an offset term of the form 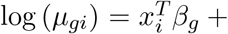 offset. By default, the offset term in *edgeR* is sample-specific and defined as log (*L*_*i*_), where *L*_*i*_ = *η*_*i*_*N*_*i*_ is the effective library size. *η*_*i*_ are sample-specific scaling factors, which are commonly computed by the TMM method. *DESeq2* uses sample-specific offset terms of the form log (*s*_*i*_) by default, where the *s*_*i*_ are size factors computed with the median ratio method. Importantly, both *DESeq2* and *edgeR* methods allow users to specify a custom offset matrix that can account for sample- and gene-specific effects at the count level.

### Gene-specific exponent corrected normalization

*gecco* obtains library size-normalized counts for downstream analyses by regressing out gene-specific library size effects estimated from the mean model. Specifically, *gecco* fits an NB-GLM with gene-specific coefficients *γ*_*g*_ of the form

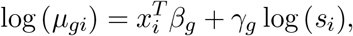

to account for gene-specific library size effects. The estimates 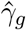 can then be used to compute observation-specific normalization factors of the form 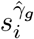 yielding normalized counts 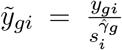 This formulation generalizes the count scaling approach of *DESeq2*’s *counts* method, which divides gene counts by the size factor *s*_*i*_, by replacing it with the gene-specific exponent-corrected quantity 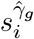 Our approach makes use of *DESeq2*’s median ratio method to estimate *s*_*i*_, although the framework is general and can be applied to the *edgeR* modeling framework as well.

### Usage and implementation

The R function *gecco* was implemented to compute library size-normalized RNA-seq counts. Given a matrix of raw counts *y*_*gi*_, *gecco* calculates size factors and fits gene-wise NB-GLMs to estimate gene-level coefficients *γ*_*g*_ corresponding to the log-size factor term log(*s*_*i*_), while optionally adjusting for sample-specific covariates *x*_*i*_. Size factors are calculated via the median ratio method, using the *estimateSizeFactorsForMatrix* function in *DESeq2*. Gene-wise NB dispersions and model coefficients are estimated using *edgeR*’s *estimateDisp* and *glmFit* functions, which implement empirical Bayes dispersion estimation and GLM fitting, respectively. Normalized counts are then computed by dividing the raw counts *y*_*gi*_ by 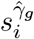.

We note that *gecco* is not intended for computing normalized counts that are then subsequently used in DE analysis with either *DESeq2* or *edgeR*. These DE methods require raw count matrices to properly model the mean-variance relationship under the NB modeling framework, as substituting normalized counts would violate the underlying statistical assumptions and ignore uncertainty in parameter estimation. Instead, our normalization strategy can be incorporated into the DE modeling framework by modifying how library size effects are handled in each method. Rather than treating size factors as fixed offsets, we include the log-size factors log(*s*_*i*_) as an additional term in the design matrix, leading to the mean model log 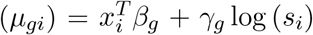. In *DESeq2*, we first estimate sample-specific size factors *s*_*i*_ using the median ratio method, set the normalization factors (which default to *s*_*i*_) to one for all samples so that no fixed offset is applied, and include log(*s*_*i*_) as an explicit covariate in the design formula, allowing *γ*_*g*_ to be estimated for each gene. An analogous approach is applied in *edgeR* by including log-effective library sizes as an explicit covariate in the design formula and setting an all-zero offset matrix in the GLM functions of *edgeR*.

In this article, we refer to the integration of *gecco* within the *DESeq2* and *edgeR* modeling frameworks as *gecco DESeq2* and *gecco edgeR*, respectively. Detailed usage guidelines and function code are available on GitHub at https://github.com/RUY28/LibrarySize-code.

### RNA-seq profiling in large-scale experiments

To assess the extent to which different normalization strategies remove residual library size effects from real RNA-seq data, we analyzed 105 publicly available human RNA-seq datasets obtained from the Gene Expression Omnibus (GEO) and The Cancer Genome Atlas (TCGA). For GEO, we selected RNA-seq datasets with samples annotated as *Homo sapiens*, sequenced on Illumina platforms, and containing between 50 and 300 samples, for which raw gene-level count matrices were available. For TCGA, we obtained raw gene-level count matrices from the TCGA Pan-Cancer dataset (GSM1536837), comprising a total of 9,264 tumor samples across 24 cancer types. The GEO series number from each dataset is provided in Supplementary Table S1.

To ensure consistent analyses across datasets, we constructed a reference gene list based on the *Homo sapiens* genome annotation GRCh38. Gene annotation was obtained from the Gene Transfer Format (GTF) file downloaded from the Ensembl database version 108. From this GTF file, we retained gene features annotated with the ensembl havana source and classified as protein-coding or long non-coding RNA (lncRNA) genes. After filtering, 19,068 genes were retained.

For datasets with Ensembl gene identifiers, raw count matrices were directly subset to include only genes present in the reference gene list. For datasets annotated using gene symbols, gene identifiers were first mapped to Ensembl IDs, and only genes with successful mappings were retained for subsequent analyses. For datasets lacking gene annotations, we instead applied an expression-based filtering strategy to remove lowly expressed genes. Specifically, counts were transformed to log-CPM using `edgeR::cpm`, and genes with mean log-CPM below the median expression level across all genes were excluded. This procedure ensured that downstream analyses were restricted to informative gene sets.

For each filtered dataset, we applied multiple widely used normalization methods to assess residual library size effects on the resulting counts. In our study we included the median ratio normalization implemented in *DESeq2*, CPM, computed in *edgeR* with both raw library size (CPM_LS) and effective library size (CPM_ELS), conditional quantile normalization (CQN), EDASeq with both GC-content normalization (EDA_GC) and gene-length normalization (EDA length), and our proposed *gecco* normalization. Specifically, median ratio normalized counts were obtained using the *counts* method from *DESeq2* with argument *normalized=TRUE*, which divides raw counts by the estimated size factors *s*_*i*_. CPM values were obtained with the *cpm* function from *edgeR*. For *edgeR*, CPM values using effective library size were computed by estimating TMM scaling factors with *normLibSizes* prior to calling *cpm*. CQN normalization was applied by jointly adjusting for GC content, gene length, and library size, yielding log-transformed normalized counts through the sum of the fitted values and offsets returned by the *CQN* function. For EDASeq-based normalization, within-lane normalization was performed separately for GC content or gene length using loess smoothing, followed by between-lane normalization using the upper-quartile method, and normalized counts were extracted using *normCounts*. Our proposed *gecco* normalized counts were obtained using the *gecco* function.

We assessed the residual dependence of normalized counts 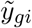 on library size by fitting gene-wise linear models of the form 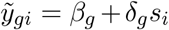. We evaluated the significance of the coefficient *δ*_*g*_, which represents the association between normalized counts and library size, using a two-sided t-test. P-values were adjusted for multiple testing using the Benjamini-Hochberg procedure [22]. For each dataset and normalization method, we summarized the results as the proportion of genes with BH-adjusted *p <* 0.05, computed as the number of significantly associated genes divided by the total number of filtered genes.

### Simulated datasets

We designed simulation studies to assess how read count normalization, DE analysis, and rhythmicity analysis are affected by the library size correction approach presented in this paper. Datasets were simulated under two scenarios differing in the specification of the mean model.

In the first scenario, data were generated to emulate our in-house human anterior cingulate cortex (ACC) RNA-seq dataset, comprising 239 samples and 16,526 filtered genes, with 119 samples from layer III and 120 samples from layer V (referred to as the in-house1 dataset in Supplementary Table S2). Specifically, the mean model had the form log (*µ*_*gi*_) = *β*_0,*g*_ +*β*_1,*g*_*x*_*i*_ +*γ*_*g*_ log (*s*_*i*_), where the binary variable *x*_*i*_ denoted the brain layer (layer III and layer V), and *s*_*i*_, estimated from the raw in-house count data, denoted the sample-specific size factor (Supplementary Fig. S1). The gene-wise intercepts *β*_0,*g*_ were sampled from a normal distribution with mean 2 and standard deviation 0.5. Simulation scenarios varied in the degree of differential expression *β*_1,*g*_ and the gene-specific library size effect *γ*_*g*_. Differential expression was generated by setting *β*_1,*g*_ to values of *±*0.2, *±*0.3, and *±*0.4, separately for each scenario, to establish balanced 500 up- and 500 down-regulation across 1,000 genes. Gene-specific coefficients *γ*_*g*_ were sampled from a normal distribution with mean 1 and standard deviation ranging from 0 (null model without gene-specific library size effect) to 0.8 in each scenario. The maximum standard deviation value was chosen to reflect the empirical variability observed in estimated *γ*_*g*_ values from the in-house ACC dataset (Supplementary Fig. S2). *γ*_*g*_ values were estimated following the procedure described in the Methods section, wherein the offset is set to zero in *edgeR* and the model is fitted using *glmFit*. Counts were simulated following an NB model with mean *µ*_*gi*_ and gene-specific dispersions *α*_*g*_, which were estimated from the in-house ACC dataset using *DESeq2* and used as fixed parameters in the simulation. (Supplementary Fig. S3). A total of 50 datasets were generated for each simulation scenario.

In the second scenario, data were generated using a cosinor mean model, as typically used in circadian rhythmicity analyses. Specifically, the mean model had the form log (*µ*_*gi*_) = *M* + *A*_*g*_ cos (*ω*(*t*_*i*_ − *ϕ*)) + *γ*_*g*_ log (*s*_*i*_). Under this parametrization, *M* is the mesor parameter, *A*_*g*_ is the gene-specific rhythmic amplitude, *ω* = 2*π/*24 is the circadian frequency, *ϕ* is the phase, and *t*_*i*_ denotes the sample-specific collection time point. In this simulation, we set *M* = 5, *ϕ* = 6, and values of *t*_*i*_ sampled at 50 uniformly distributed time points between 0 and 24 hours. Rhythmicity was introduced for 1,000 randomly chosen genes, for which amplitude values *A*_*g*_ were sampled from a truncated normal distribution with mean values in the set 0.5, 1.0, 1.5, each defining a separate simulation scenario, and a standard deviation of 0.5, which is a typical value setting for circadian analysis simulation [23]. Non-rhythmic genes were assigned *A*_*g*_ = 0. Size factors *s*_*i*_ and the gene-specific coefficients *γ*_*g*_ were the same as in the previous simulation scenario. Counts and dispersions were simulated as in the first scenario, with 50 datasets generated per simulation scenario.

## Results

### Normalized counts remain associated with library size in large-scale RNA-seq experiments

We fitted gene-wise linear regression models of normalized counts on the estimated size factor *s*_*i*_ to verify the existence of any residual dependence on library size after normalization. In most of the 105 RNA-seq datasets, the size factor *s*_*i*_ remained significantly associated with normalized counts even after normalization (BH-adjusted *p <* 0.05) for a substantial fraction of genes for all existing normalization methods (Fig. 1a). From the plot, we observed that none of these traditional normalization approaches consistently removed the dependence on library size. In addition, we repeated our analyses using raw library size as a covariate in the linear model and observed similar results (Supplementary Fig. S4).

**Figure 1:**
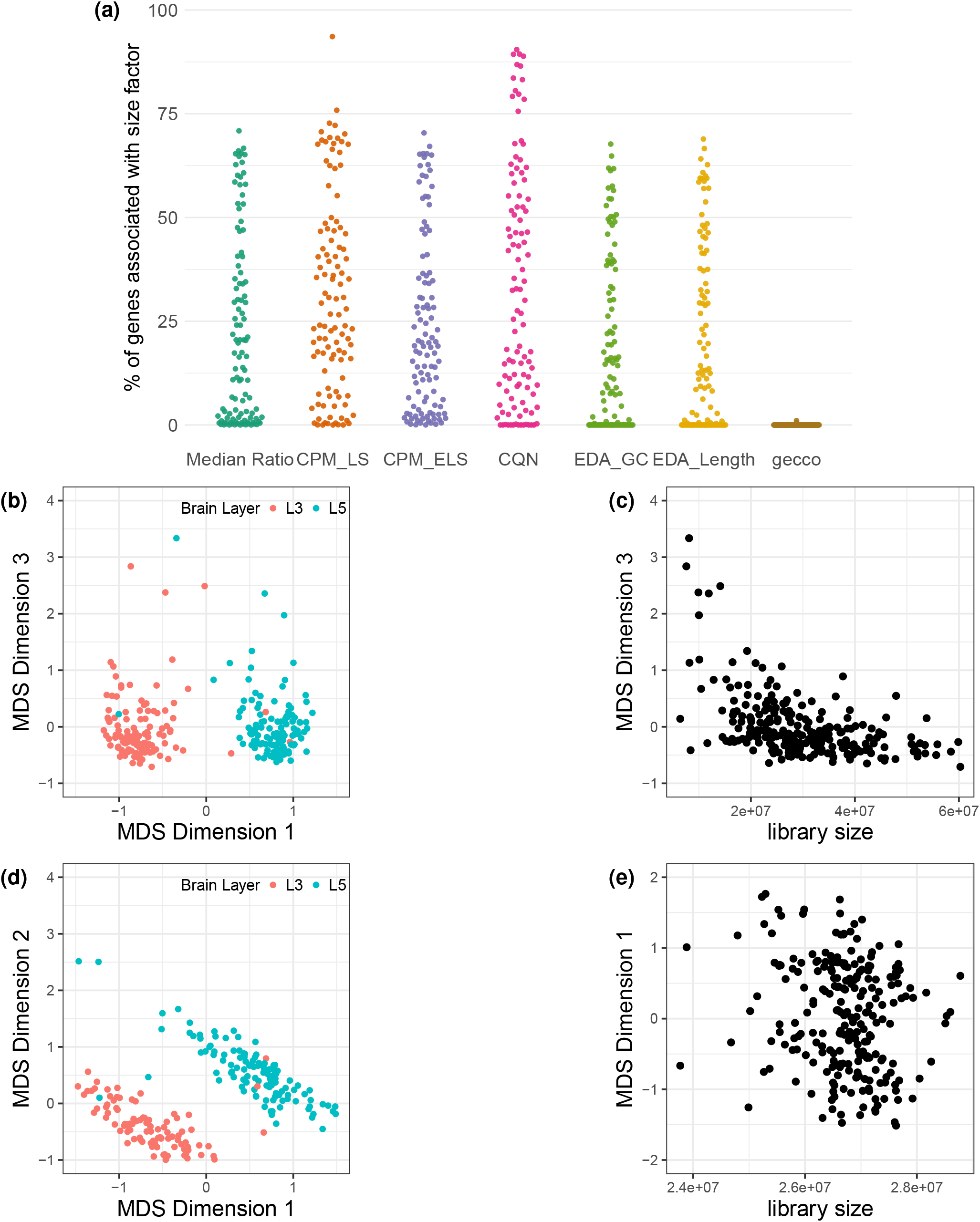
(a) Proportion of genes from each of the 105 RNA-seq datasets for which gene-wise normalized counts were significantly associated with *s*_*i*_ (BH-adjusted *p <* 0.05) under multiple normalization methods. (b) MDS plot of log-CPM from the ACC dataset, showing sample clustering by brain layer. (c) Association between MDS Dimension 3 and raw library size under CPM normalization, suggesting residual library size effects. (d) MDS plot of *gecco*-normalized counts from the ACC dataset, showing preserved sample clustering by brain layer. (e) Lack of association between MDS Dimension 1 and library size after *gecco* normalization, suggesting improved removal of library size-related technical variation.

We therefore examined whether our proposed *gecco* approach could effectively mitigate this residual effect. As shown in Fig. 1a, applying *gecco* normalization mitigated library size-associated artifacts by regressing out library size effects from the read counts. Across all datasets, the proportion of genes with significant residual association with library size (BH-adjusted *p <* 0.05) was consistently low, with a maximum of 1.05% of genes showing significant residual association.

Fig. 1b–e illustrate the effect of library size on normalized counts using multidimensional scaling (MDS) plots. Under CPM normalization (Fig. 1b–c), while samples from different brain layers of the ACC dataset were separated from one another along MDS Dimension 1, we observed an association between MDS Dimension 3 and library size, suggesting that CPM normalization does not fully eliminate library size-related variability. In contrast, *gecco* normalization (Fig. 1d–e) substantially attenuated the relationship between MDS dimensions and library size, indicating improved removal of library size effects. Importantly, samples continued to cluster according to brain layer after gene-specific exponent correction, demonstrating that *gecco* effectively mitigated technical variation while preserving biologically meaningful structure.

### Gene-specific library size effects in DE analyses of large-scale RNA-seq experiments

Next, we assessed whether this library size artifact propagates to downstream DE analysis. Differential expression was assessed by testing the coefficient associated with log(*s*_*i*_), while retaining the default log-transformed size factors as offsets and excluding any additional clinical covariates, yielding the mean model log (*µ*_*gi*_) = (1 + *γ*_*g*_) log (*s*_*i*_). We performed an analogous analysis within the *edgeR* framework by including log(*L*_*i*_) as a covariate. This analysis was equivalent to testing whether the effect of library size on read counts is uniform across genes, under which no DE genes would be expected if library size were fully accounted for by the offset. Among the 105 publicly available RNA-seq datasets, 56 had over 20% of genes significantly associated with log(*s*_*i*_) within the *DESeq2* framework, and 70 had over 20% associated with log(*L*_*i*_) within the *edgeR* framework (BH-adjusted *p <* 0.05) (Supplementary Fig. S5). These results highlight that differences in library size across RNA-seq samples still resulted in spurious DE signals even after standard normalization.

We acknowledge scenarios in which library size is correlated with a known biological covariate that is expected to introduce DE. In such scenarios, read counts would be expected to remain correlated with library size in an unadjusted analysis. We performed DE analyses on eight datasets (five in-house and three public datasets) for which complete experimental design information was available. Descriptions of the in-house datasets and GEO series numbers for the public datasets are provided in Supplementary Table S2. In these analyses, the mean model was adjusted for known clinical covariates and library size, modeled additively, either by including log(*s*_*i*_) as a covariate in the *DESeq2* model or log(*L*_*i*_) in the *edgeR* model. For example, under the *DESeq2* framework, the mean model is given by 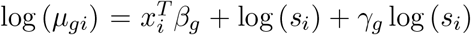. Across all eight datasets, we observed that at least 20% of genes were DE relative to library size under both the *DESeq2* and *edgeR* frameworks (Supplementary Fig. S6), suggesting that this artifact persisted even after accounting for relevant clinical covariates. These results highlight that commonly used normalization methods do not fully correct for library size effects in large-scale RNA-seq datasets.

To further assess model fit, we employed the Bayesian Information Criterion (BIC) to compare the standard *DESeq2* and *edgeR* GLMs, which include log-size factors as fixed offsets, against their *gecco*-based counterparts (*gecco DESeq2* and *gecco edgeR*), which instead include log(*s*_*i*_) or log(*L*_*i*_) as explicit gene-specific covariates while removing the offset terms. The BIC quantifies model fit while penalizing model complexity, enabling a principled comparison between the conventional DE GLM methods and the *gecco*-enhanced GLM that incorporates gene-specific exponent correction to account for technical bias. We conducted this analysis on two datasets: the in-house human ACC RNA-seq dataset comprising 239 samples and 16,526 filtered genes, and a publicly available human postmortem caudate RNA-seq dataset with 36 samples and 15,622 filtered genes [24]. Genes were classified as artifact genes or non-artifact genes depending on whether their normalized counts were significantly associated with library size upon median ratio normalization (BH-adjusted *p <* 0.05). We then compared BIC differences between each pair of models (standard GLM minus *gecco* GLM) across these two groups of genes (Fig. 2 for *DESeq2* results and Supplementary Fig. S7 for *edgeR* results). For artifact genes, the *gecco* models consistently achieved lower BIC values, indicating improved model fit. For non-artifact genes, BIC differences were close to zero, suggesting that *gecco* does not introduce overfitting when library size artifacts are absent. Together, these findings highlight the advantage of gene-specific exponent correction in capturing gene-level technical bias while preserving robust performance for well-behaved genes, improving overall model adequacy compared to the standard *DESeq2* and *edgeR* frameworks.

**Figure 2:**
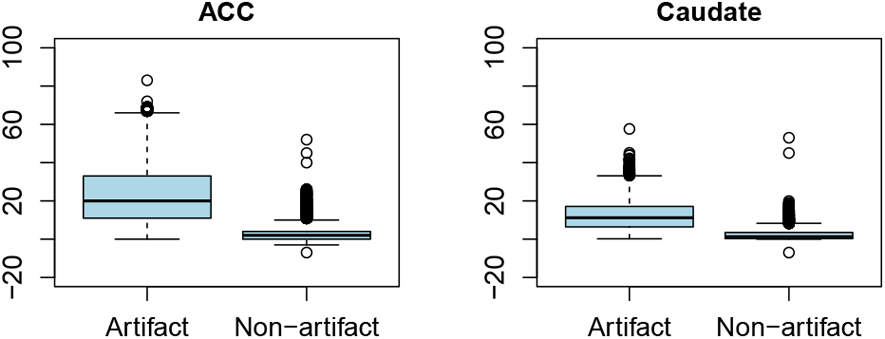
ACC and Caudate RNA-seq Data: Differences in BIC Between *DESeq2* and *gecco DESeq2* Models for artifact and Non-artifact Genes.

### gecco normalization removes library size effect from RNA-seq counts in simulations

After generating the simulated data, the primary goal was to evaluate whether our proposed method could effectively mitigate library size bias. Fig. 3 shows the proportion of genes whose normalized count levels remained significantly associated with library size (BH-adjusted *p <* 0.05) under each simulated dataset when the biological effect size *β*_1,*g*_ = *±* 0.3. This evaluation was performed across varying degrees of gene-specific library size effects, as determined by the standard deviation of *γ*_*g*_, and different normalization methods. Similar results were obtained for additional biological effect sizes (*β*_1,*g*_ = *±* 0.2 and *±* 0.4)(Supplementary Fig. S8). The results showed that when the standard deviation of *γ*_*g*_ was set to zero, which corresponds to the null scenario without gene-specific library size effects, all normalization methods successfully eliminated the library size bias. This was expected, as there was no gene-specific variation in library size effects under this scenario. As the standard deviation of *y*_*g*_ increased, the library size artifact became more pronounced. In such cases, *gecco* more effectively mitigated library size bias compared to conventional normalization approaches.

**Figure 3:**
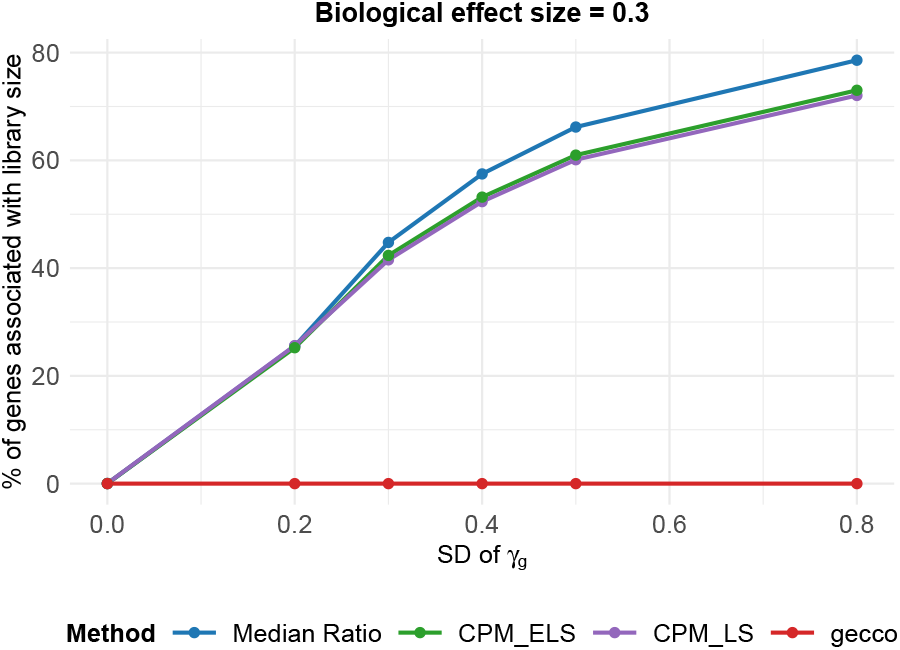
Persistence of library size-associated correlations (BH-adjusted *p <* 0.05) across normalization methods and levels of *γ*_*g*_, given a biological effect size of *β*_1,*g*_ = *±*0.3.

### Coupling gecco with DE and rhythmicity tools provides better model fit, improved power and correct FDR control in simulations

To further evaluate the impact of our method on downstream analyses, we investigated whether incorporating gene-specific exponent correction improves the detection accuracy of DE genes in simulated data. Specifically, we compared the conventional versions of *DESeq2* and *edgeR* with our proposed *gecco* approach. Across a range of library size variability (standard deviation (sd) of *γ*_*g*_ ∈ {0, 0.2, 0.3, 0.4, 0.5, 0.8}) and biological effect sizes (*β*_1*g*_ = 0.2, 0.3, 0.4), *gecco DESeq2* and *gecco edgeR* consistently outperformed their conventional counterparts *DESeq2* and *edgeR* in DE gene detection (Fig. 4).

**Figure 4:**
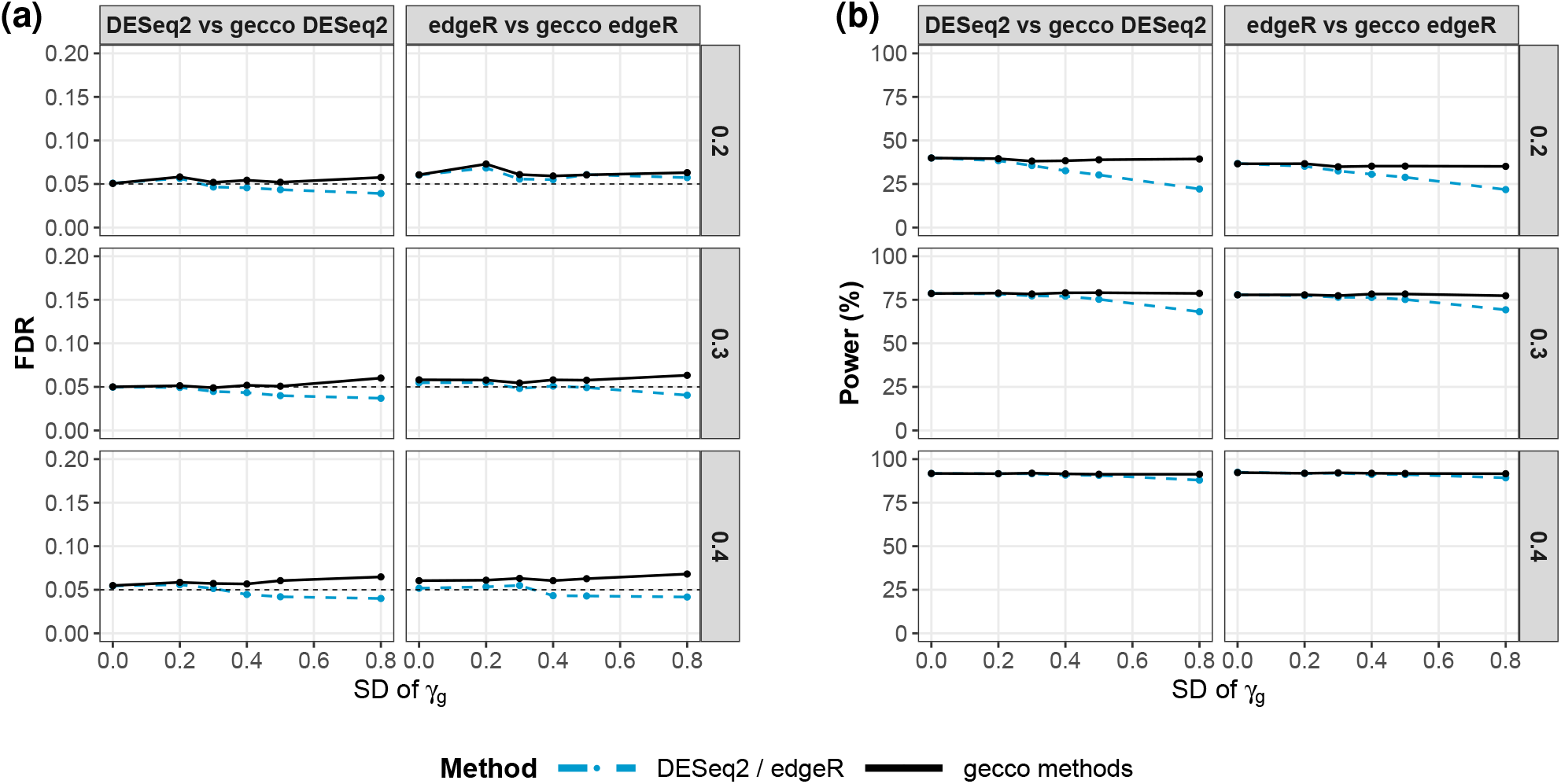
Differential expression analysis performance comparison across varying standard deviations of *γ*_*g*_ (0, 0.2, 0.3, 0.4, 0.5, and 0.8). Rows correspond to biological effect sizes (*β*_1*g*_ ∈ {0.2, 0.3, 0.4}) and columns correspond to comparisons between *DESeq2* and *edgeR* and their respective *gecco* counterparts. Panel (a) shows the observed FDR. Panel (b) shows the observed statistical power. The dashed line in indicates FDR = 0.05. Reported observed FDR and statistical power values are averaged across 50 simulation replicates.

As shown in Fig. 4a, the *gecco*-based methods maintained observed false discovery rates (FDRs) close to the nominal level of 0.05 across all settings. In addition, as shown in Fig. 4b, *gecco* achieved comparable or higher statistical power across all scenarios. When the sd of *γ*_*g*_ = 0, both approaches performed nearly identically, as expected in the absence of gene-specific library size variation. As the variability of *γ*_*g*_ increased, the advantage of *gecco* became more substantial, particularly at smaller effect sizes, whereas for larger *β*_1*g*_ the dominant biological signal reduced the relative impact of library size bias.

To further evaluate the FDR control of *DESeq2* and *edgeR* coupled with *gecco*, we also performed a gene ranking analysis to assess the number of false discoveries in the set of top-ranked most significant genes from each method (Supplementary Fig. S9 and Fig. S10). Overall, the analysis shows that both *gecco*-based methods provided the smallest number of false discoveries across all combinations of library size artifact severity and biological effect size. Collectively, these results demonstrate that incorporating gene-specific exponent correction effectively mitigates library size artifacts and improves both the statistical power and FDR control in downstream DE gene detection.

We next evaluated whether incorporating *gecco* normalization improves detection accuracy in rhythmicity analysis using simulated data. Unlike DE analysis where library size was modeled as a gene-specific covariate within existing *DESeq2* and *edgeR* pipelines, rhythmicity pipelines such as DiffCirca [6] typically take normalized counts as input. In this context, we applied our *gecco* normalization prior to rhythmicity analysis to assess its impact. We compared four normalization methods applied to the same simulated data: median ratio, CPM LS, CPM ELS, and *gecco*. Across a range of library size variability (sd of *γ*_*g*_ ∈ {0, 0.2, 0.3, 0.4, 0.5, 0.8}) and rhythmic signal strengths *A*_*g*_ ∈ {0.5, 1, 1.5}, *gecco* normalization consistently improved the accuracy of circadian gene detection compared to other conventional normalization methods (Fig. 5). As shown in Fig. 5a, *gecco* maintained FDRs close to the nominal level of 0.05 across all settings, whereas alternative normalization methods, particularly CPM_LS, exhibited substantial inflation of FDR as library size variability increased. Median ratio and CPM_ELS showed moderate deviations but remained less stable than *gecco* under increasing bias. Meanwhile, Fig. 5b shows that *gecco* achieved comparable or better statistical power to existing methods across all scenarios, indicating that improved error control was not achieved at the expense of sensitivity. Consistent with the DE analysis results, the advantage of *gecco* became more pronounced as the variability of *γ*_*g*_ increased, particularly under weaker rhythmic signals (*A*_*g*_ = 0.5).

**Figure 5:**
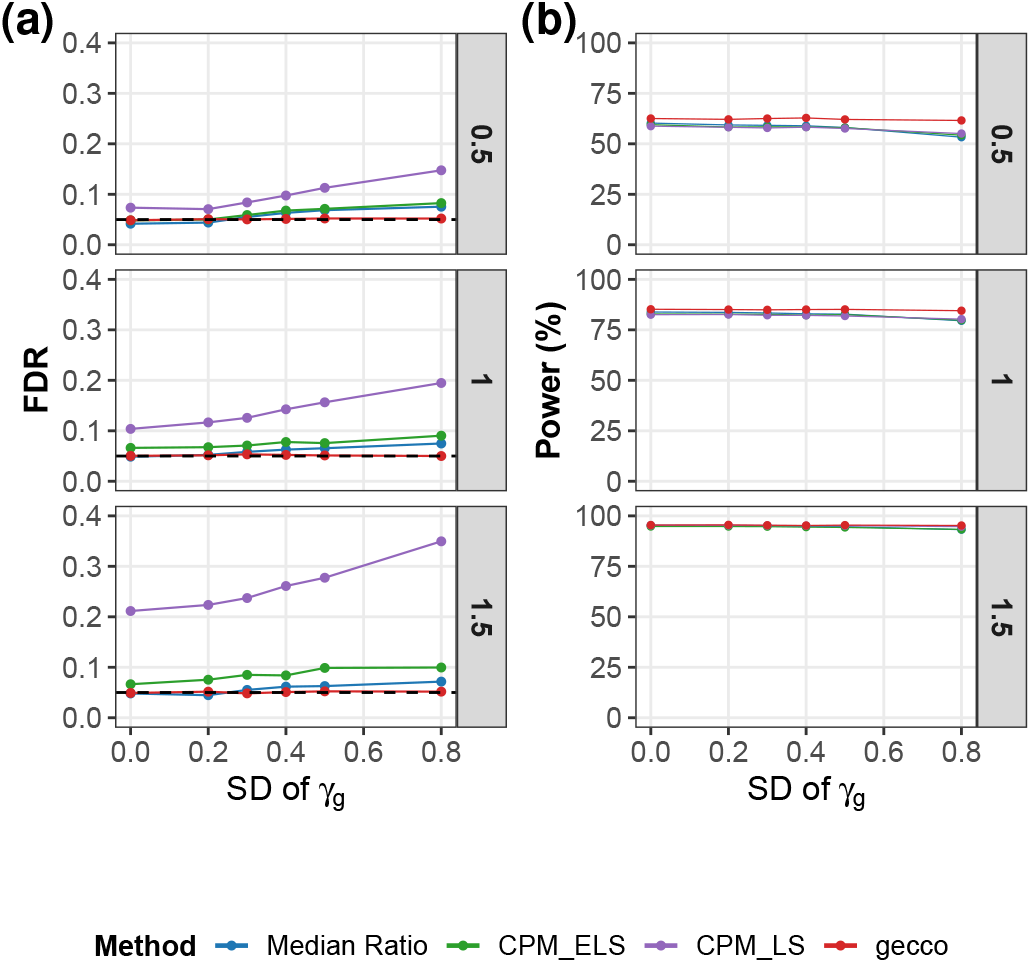
Rhythmicity performance comparison across varying standard deviations of *γ*_*g*_ (0, 0.2, 0.3, 0.4, 0.5, and 0.8), with rows representing rhythmic signal strengths *A*_*g*_ (0.5, 1, and 1.5), comparing median ratio, CPM_LS, CPM_ELS, and *gecco* normalization methods. Panel (a) shows the observed FDR. Panel (b) shows the observed statistical power. The dashed line in (a) indicates FDR = 0.05. Reported observed FDR and statistical power values are averaged across 50 simulation replicates.

Supplementary Fig. S11 presents the cumulative number of false discoveries among the top-ranked rhythmic genes under each normalization method, providing a complementary ranking-based perspective on FDR control. *gecco* consistently accumulated fewer false discoveries than competing normalization approaches across most settings, with the largest improvements observed under higher library size variability and weaker rhythmic signals. Taken together, these results support the use of *gecco* normalization as a more reliable preprocessing step for rhythmicity analysis in the presence of library size variability.

### Enhanced downstream analytical performance on real data: Differential expression analysis

Here we present a real data application to more comprehensively evaluate the effect of library size and our proposed *gecco* approach in DE analysis. We performed a DE analysis using *gecco DESeq2* and compared the results with those obtained from the conventional *DESeq2* pipeline, as described in the Methods section. We applied both methods to the in-house human postmortem dorsolateral prefrontal cortex (DLPFC) striatum RNA-seq dataset consisting of 249 samples and 13,369 filtered genes (referred to as the in-house5 dataset in Supplementary Table S2). We tested for DE between Major Depressive Disorder (MDD) cases and controls, adjusting for sex and age as covariates. Genes with Benjamini-Hochberg adjusted p-values below 0.05 were declared DE.

The resulting Venn diagram (Supplementary Fig. S12) illustrates the overlap between DE genes identified by *DESeq2* and *gecco DESeq2*. Of the DE genes identified, 41 were detected by both methods, 11 were unique to *gecco DESeq2*, and 1 was unique to *DESeq2*. In the absence of ground truth for DE genes, we further evaluated the biological relevance of genes found to be DE by each method using pathway enrichment analysis.

Fig. 6 shows pathways enriched among DE genes identified by either *DESeq2* or *gecco_DESeq2*. Compared to *DESeq2*, DE genes identified by *gecco_DESeq2* not only recovered established MDD-related pathways from the literature, including the TNF signaling pathway [25], NF-*κ*B signaling pathway [26], NOD-like receptor signaling pathway [27], and PI3K-AKT signaling pathway [28], but also identified additional MDD-relevant pathways, including Herpes simplex virus 1 infection [29], MAPK signaling pathway [30–32], ECM-receptor interaction [33], and IL-17 signaling pathway [34]. Moreover, enrichment signals for shared pathways were generally stronger under the *gecco*-based analysis, indicating improved sensitivity.

**Figure 6:**
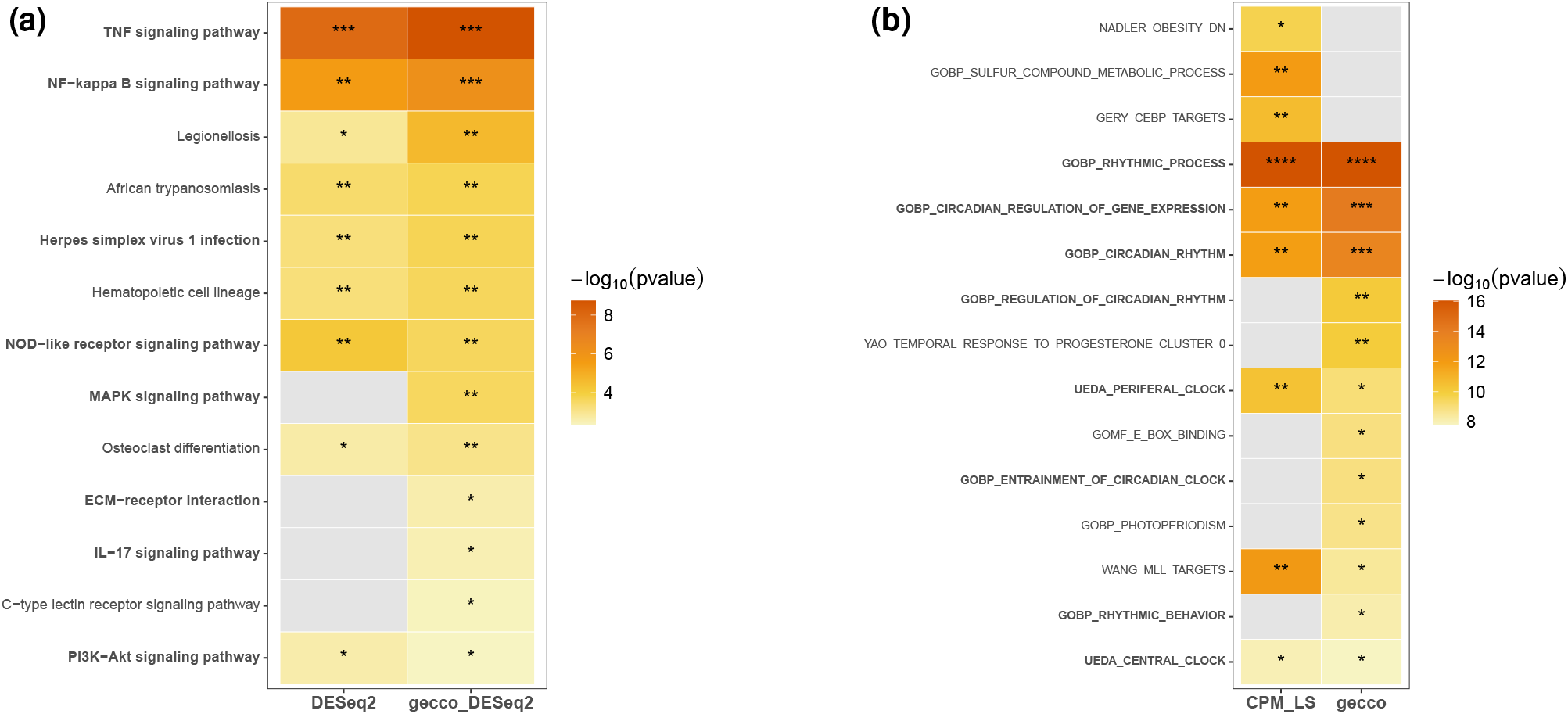
(a) Heatmap of pathways enriched among DE genes identified by *DESeq2* and *gecco DESeq2*. Color indicates −log_10_(*p*-value), and asterisks denote significance levels (* *p <* 10^−1^, ** *p <* 10^−3^, *** *p <* 10^−6^, **** *p <* 10^−9^). Pathways in **bold** have been previously associated with MDD. (b) Heatmap of pathways enriched among circadian genes identified by DiffCirca on CPM_LS and *gecco* normalized datasets. Color indicates −log_10_(*p*-value), and asterisks denote significance levels (* *p <* 10^−7^, ** *p <* 10^−10^, *** *p <* 10^−13^, **** *p <* 10^−16^). Pathways in **bold** have been previously associated with circadian regulation.

### Enhanced downstream analytical performance on real data: Rhythmicity analysis

To evaluate the impact of library size artifacts on rhythmicity detection in real RNA-seq data, we applied *gecco* normalization to a circadian time-course dataset (GSE248462). This dataset profiles genome-wide transcriptional dynamics across circadian time in mouse tissues, including the arcuate nucleus and left nodose ganglia. The dataset consists of samples collected at six circadian time points (0–24 hours at 6-hour intervals), with 6 biological replicates per time point, each derived from independent tissue dissections.

Raw read counts were obtained from GEO and restricted to the 24 control samples. Genes with low expression were filtered, resulting in 12,614 genes retained for downstream analyses. Two normalization strategies were then applied: standard log_2_-CPM normalization (referred to as CPM_LS above) and *gecco* normalization. To quantify residual library size dependence after normalization, we assessed the association between normalized gene counts and library size. Under log_2_CPM normalization, 2,581 genes remained significantly associated with library size (BH-adjusted p *<* 0.05), indicating substantial residual library size bias. In contrast, after applying *gecco* normalization, no genes showed significant association with library size, demonstrating that *gecco* effectively removes residual library size effects. Circadian rhythmicity was then assessed using the *DCP Rhythmicity* function from the Diffcirca framework, with rhythmic genes defined as those with BH-adjusted *p <* 0.05. Using log_2_-CPM-normalized data, 134 genes were identified as rhythmic, whereas *gecco* normalization identified 77 rhythmic genes. Of these, 59 genes were detected by both methods (see Venn diagram in Supplementary Fig. S13).

To evaluate the biological relevance of these rhythmic gene sets, pathway enrichment analysis was performed using gene sets from the Molecular Signatures Database (MSigDB) and the `clusterProfiler` framework after mapping Ensembl gene identifiers to Entrez IDs. As shown in Fig. 6b, pathways identified based on *gecco* normalization exhibited stronger and more consistent enrichment of circadian-related processes (highlighted in bold) compared to those obtained using CPM normalization. Core circadian pathways, including circadian regulation of gene expression and circadian rhythm, were robustly detected under both normalization methods. However, *gecco* generally yielded higher enrichment significance and additionally captured related pathways such as regulation of circadian rhythm. In contrast, *CPM_LS* identified additional pathways unrelated to circadian rhythm. These results suggest that *gecco* normalization enhances the recovery of circadian signals and improves the biological interpretability of rhythmicity analysis by mitigating library size-related artifacts. We further evaluated normalization performance by examining whether detected circadian genes recovered known core clock components. Of the 134 genes detected by *CPM_LS*, eight core clock genes were identified: *Arntl, Per1, Per2, Per3, Cry1, Cry2, Nr1d1*, and *Nr1d2*. In contrast, although *gecco* detected fewer genes overall (77 genes), the *gecco*-identified gene set retained all eight core clock genes and additionally identified *Clock*, a key regulator of the circadian machinery. Although *Clock* is often reported to exhibit weak or absent rhythmicity in the suprachiasmatic nucleus (SCN) [35], previous studies have shown that *Clock* can display rhythmic expression in other brain regions and peripheral tissues [36]. Fig. 7 shows the temporal expression of *Clock* under the two normalization strategies. While *CPM_LS* yielded a weak and non-rhythmic pattern (p = 0.099, *R*^2^ = 0.17), with a relatively small oscillation amplitude compared with the MESOR (*A/M* = 0.03), *gecco* normalization revealed a clear oscillatory trend with improved statistical significance and model fit (p = 0.021, *R*^2^ = 0.29), as well as a substantially larger relative amplitude (*A/M* = 0.13).

**Figure 7:**
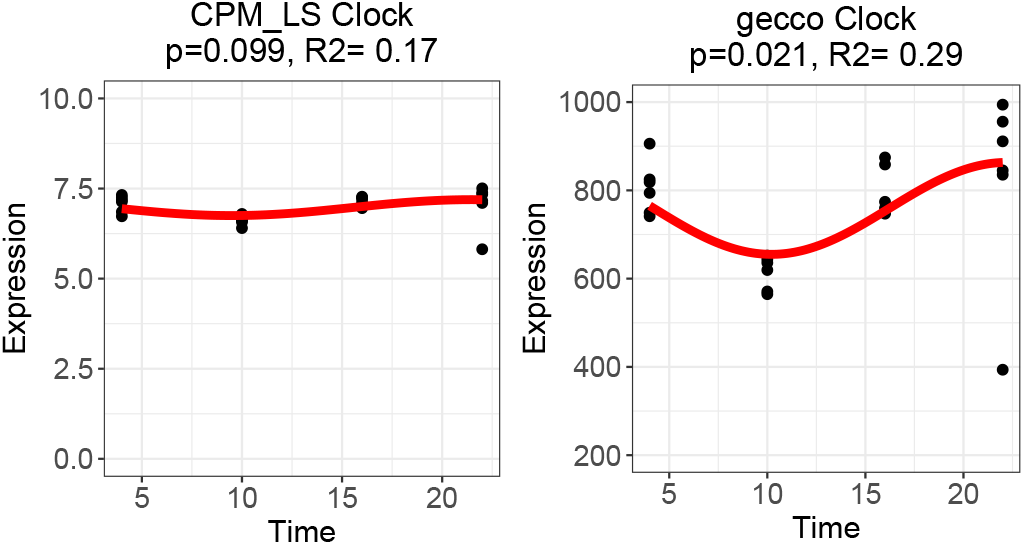
Scatterplots of gene expression across circadian time for *CLOCK* under CPM and *gecco* normalization. The fitted curve (red) illustrates the temporal trend.

Together, these results indicate that *gecco* normalization improves the recovery of biologically meaningful circadian signals and the detection of core clock genes that are missed under conventional normalization.

## Discussion

In this paper, we present a flexible statistical framework for library size normalization of bulk RNA-seq data. Normalization is a critical step in RNA-seq analysis, as it removes technical variability attributable to library size differences, enabling accurate downstream analyses such as visualization, clustering, differential expression (DE), and rhythmicity analysis. Unlike conventional approaches that apply a uniform scaling factor across all genes, our proposed method, *gecco*, addresses gene-level effects of library size that conventional methods fail to capture. This flexible formulation yields normalised counts that are effectively free of residual library size effects.

Our evaluations, based on both simulated and real datasets, demonstrate that *gecco* outperforms widely used normalization techniques in the presence of library size artifacts, mitigating library size-related artifacts and improving detection accuracy in DE and rhythmicity analyses. Importantly, in null scenarios where no gene-specific library size effects are present, our method performs comparably to existing approaches, indicating that *gecco* does not introduce overfitting or spurious signals.

Accurate normalization of library size variation is essential for ensuring valid inference in RNA-seq studies. Traditional approaches assume that library size affects all genes proportionally. By introducing a gene-specific library-size correction, our framework more effectively removes heterogeneous library size-related artifacts, resulting in expression estimates that better reflect underlying biological variation. In DE analysis, correcting gene-level sensitivity to library size reduces spurious signals driven by technical heterogeneity and yields more stable effect-size estimates. Similarly, circadian analyses, which rely on precise quantification of rhythmic amplitude and phase, are highly susceptible to subtle normalization biases. Library size correction reduces technically induced distortions in rhythmic patterns, leading to more reliable identification of rhythmic genes and more accurate phase estimation across samples or conditions. In both case studies, *gecco* yielded more biologically interpretable pathway enrichment results, recovering disease-relevant pathways in the DE analysis and circadian-related pathways in the rhythmicity analysis. Overall, gene-specific correction of library-size effects enhances the robustness and biological interpretability of multiple downstream analyses by minimizing technical confounding.

Despite these promising results, the underlying sources contributing to the observed artifacts remain unclear. We explored potential associations with known genomic features such as GC content and gene length. However, these analyses did not reveal significant associations. While it is unclear why such an effect is observed in large-scale RNA-seq datasets, we speculate that it may arise when differences in cell type composition across samples confound the relationship between library size and gene expression. In bulk RNA-seq experiments, gene expression is expected to reflect the cell type composition of each sample, and the number of sequenced RNA fragments contributed by each cell type scales with the overall library size of the sample. As a result, library size would be expected to be associated with cell type-specific genes in such scenario. Identifying the origins of this artifact remains an open question. Future work will focus on systematically investigating additional genomic and experimental variables that may contribute to this phenomenon. Such investigations may further refine normalization strategies and deepen our understanding of technical variability in RNA-seq data.

## Competing interests

No competing interest is declared.

## Author contributions statement

M.L.S, C.A.M., P.L.B. and G.C.T. conceived the study. R.Y. and W.Z. curated the data. R.Y. and D.L. performed the formal analysis. M.L.S., C.A.M. and G.C.T. acquired funding. P.L.B. and G.C.T. conducted the investigation. G.C.T. administered the project and supervised the study. K.D.K., M.L.S., C.A.M. and G.C.T. provided data and computing resources. R.Y. wrote the original manuscript draft. All authors reviewed and approved the manuscript.

## Key Points

- Conventional approaches to adjust for library size differences in large-scale RNA-seq experimental data frequently fail to fully correct for library size effects. Conventional library size adjustment methods assume that library size affects all genes proportionally, an assumption that is often violated in large-scale RNA-seq experimental data.
- *gecco* addresses the limitation of conventional library size adjustment strategies by incorporating library size as a gene-specific effect in a negative binomial generalized linear model, yielding normalized counts and downstream differential analyses that are effectively free of residual library size effects.
- *gecco* improves detection accuracy and pathway enrichment interpretability in downstream DE and rhythmicity analyses without compromising false discovery rate control or statistical power in the absence of library size artifacts.
- *gecco* is fully compatible with widely used differential gene expression analysis methods such as *DESeq2* and *edgeR*.

## Acknowledgments

This work is supported in part by funds from the National Institutes of Health [R01LM014142 and R01CA285337 to G.C.T.; P50DA046346 and R01MH111601 to R.Y., W.Z., C.A.M., M.L.S. and G.C.T.]. This study used the University of Pittsburgh Center for Research Computing and Data, RRID:SCR 022735, through the resources provided. Specifically, this work used the HTC cluster, which is supported by NIH award number S10OD028483.

## Data availability statement

The *gecco* function and analysis code are available on GitHub at https://github.com/RUY28/LibrarySize-code. Publicly available RNA-seq datasets analyzed in this study were obtained from the Gene Expression Omnibus (GEO) at https://www.ncbi.nlm.nih.gov/geo/ and The Cancer Genome Atlas (TCGA) at https://www.cancer.gov/tcga. GEO series numbers and TCGA cancer types are provided in Supplementary Tables S1 and S2.

## Notes

### Competing Interest Statement

The authors have declared no competing interest.

