## supplemental files for "Gene-specific exponent-corrected normalization for library size in bulk RNA-seq"

### Supplementary materials for "Gene-specific exponent corrected normalization on sequencing depth for bulk RNA-seq"

8 April 2026

#### Contents

**1 Supplementary Figures** **2**

**2 Supplementary Table1** **13**

**3 Supplementary Table2** **15**

[1]Department of Biostatistics and Health Data Science, University of Pittsburgh, 130 De Soto Street, Pittsburgh, PA 15261, United States,
[2]Department of Psychiatry, University of Pittsburgh, 450 Technology Drive, Pittsburgh, PA 15219, United States,

### 1 Supplementary Figures

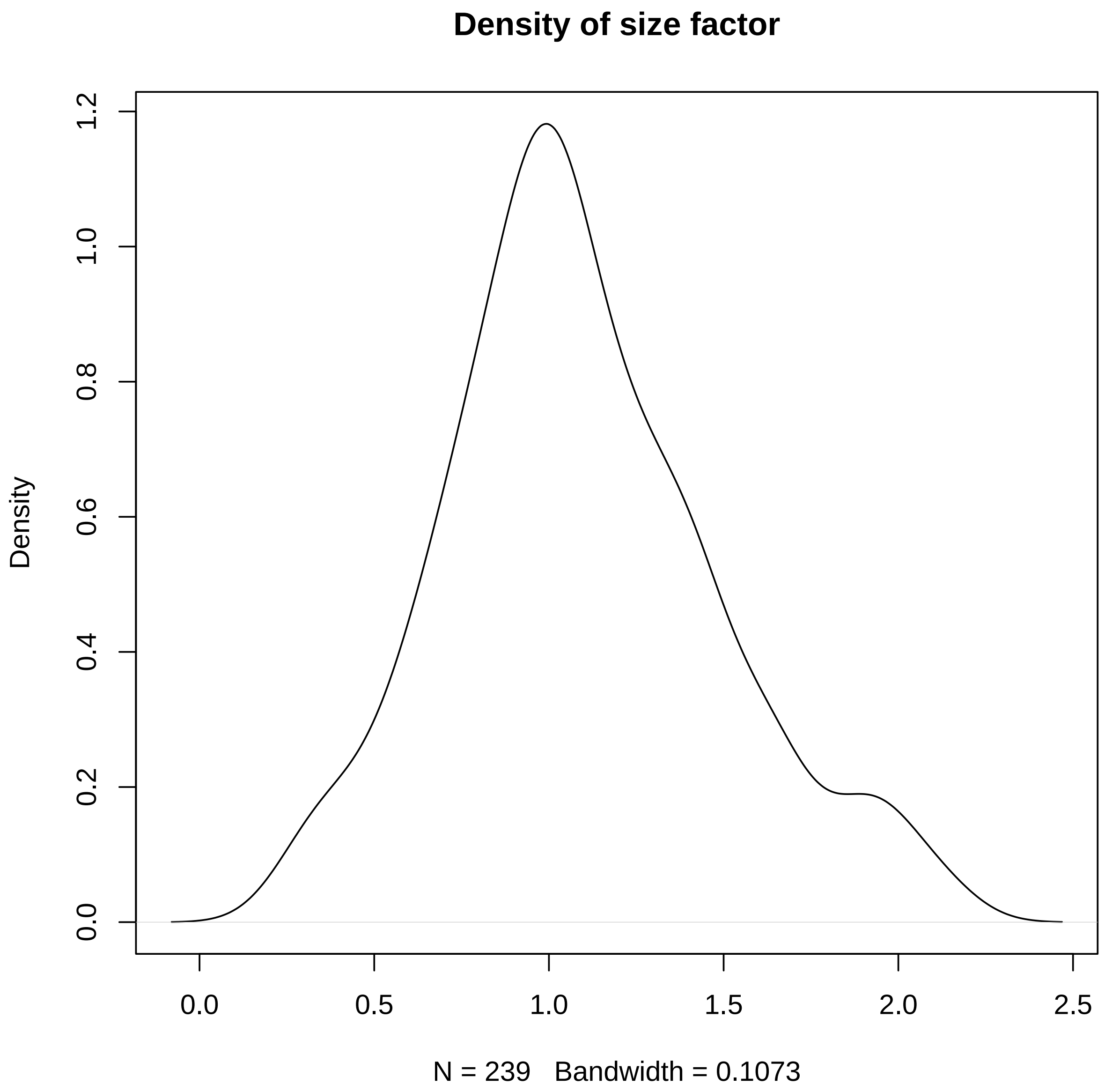


Supplementary Figure S1: Distribution of size factors $s_i$ estimated from the in-house DLPFC dataset and used in the simulation study.

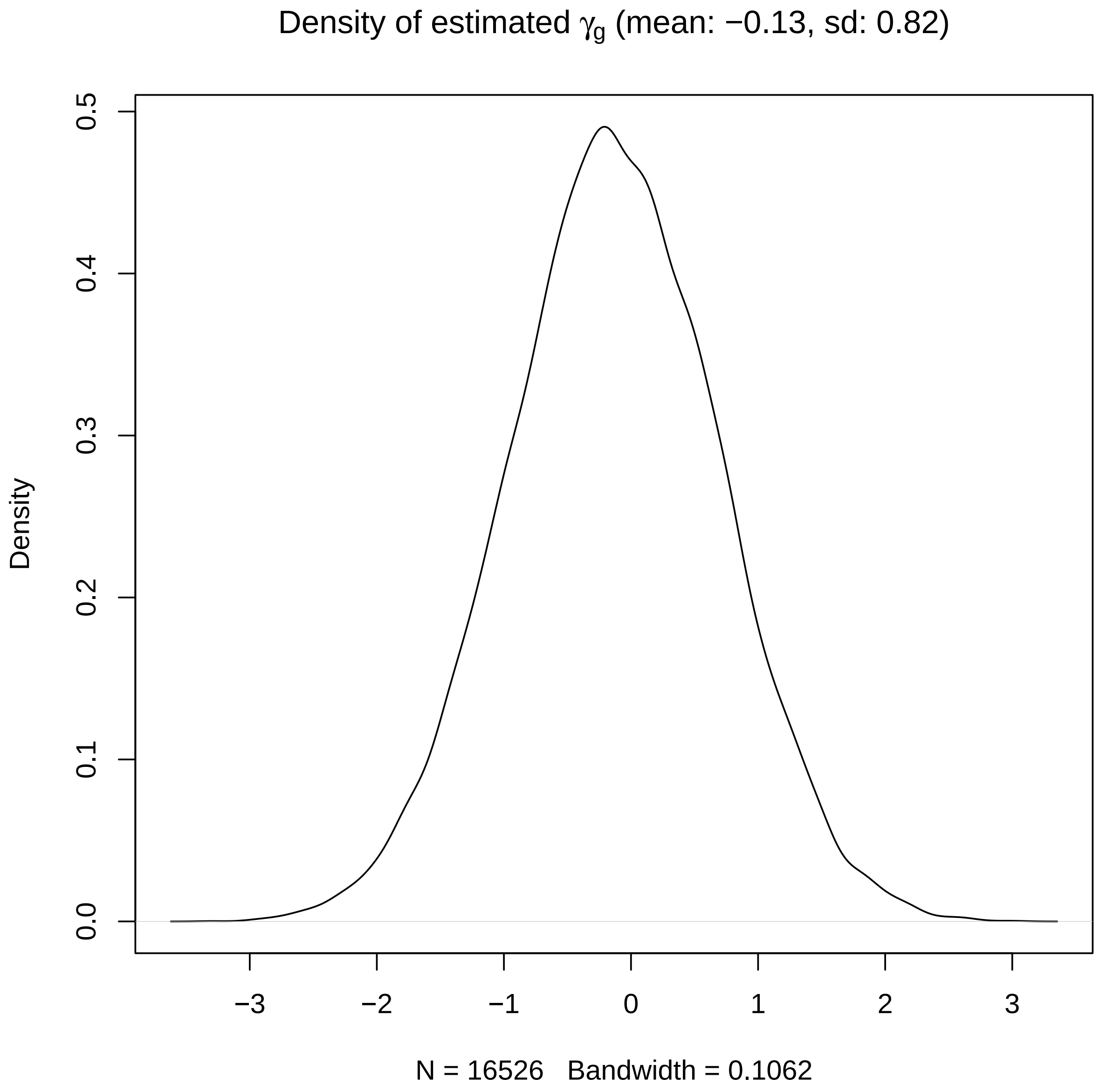


Supplementary Figure S2: Distribution of estimated $\gamma_g$ values from the in-house DLPFC dataset and used in the simulation study.

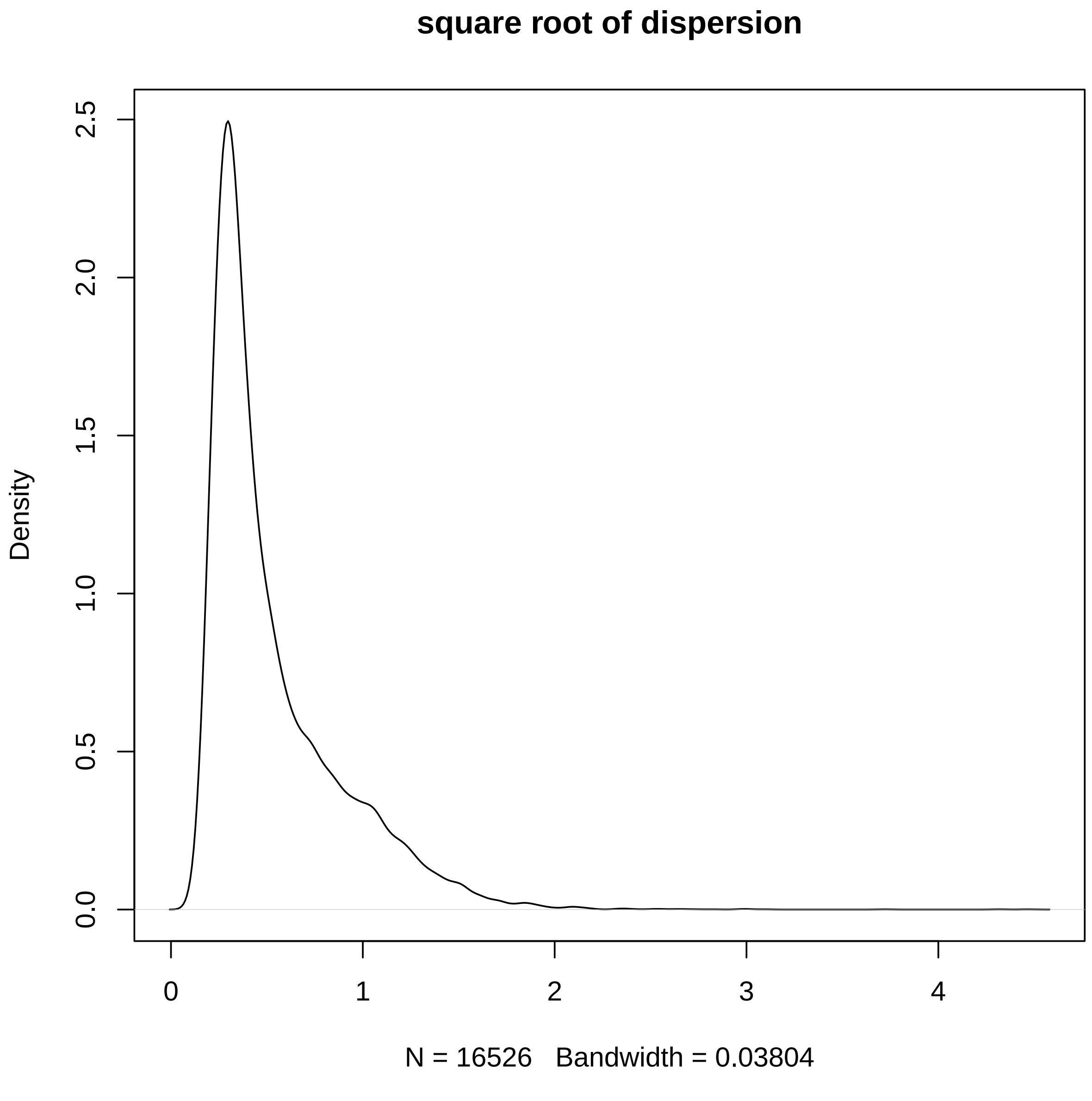


Supplementary Figure S3: Distribution of the square root of dispersion parameters estimated from the in-house DLPFC dataset and used in the simulation study.

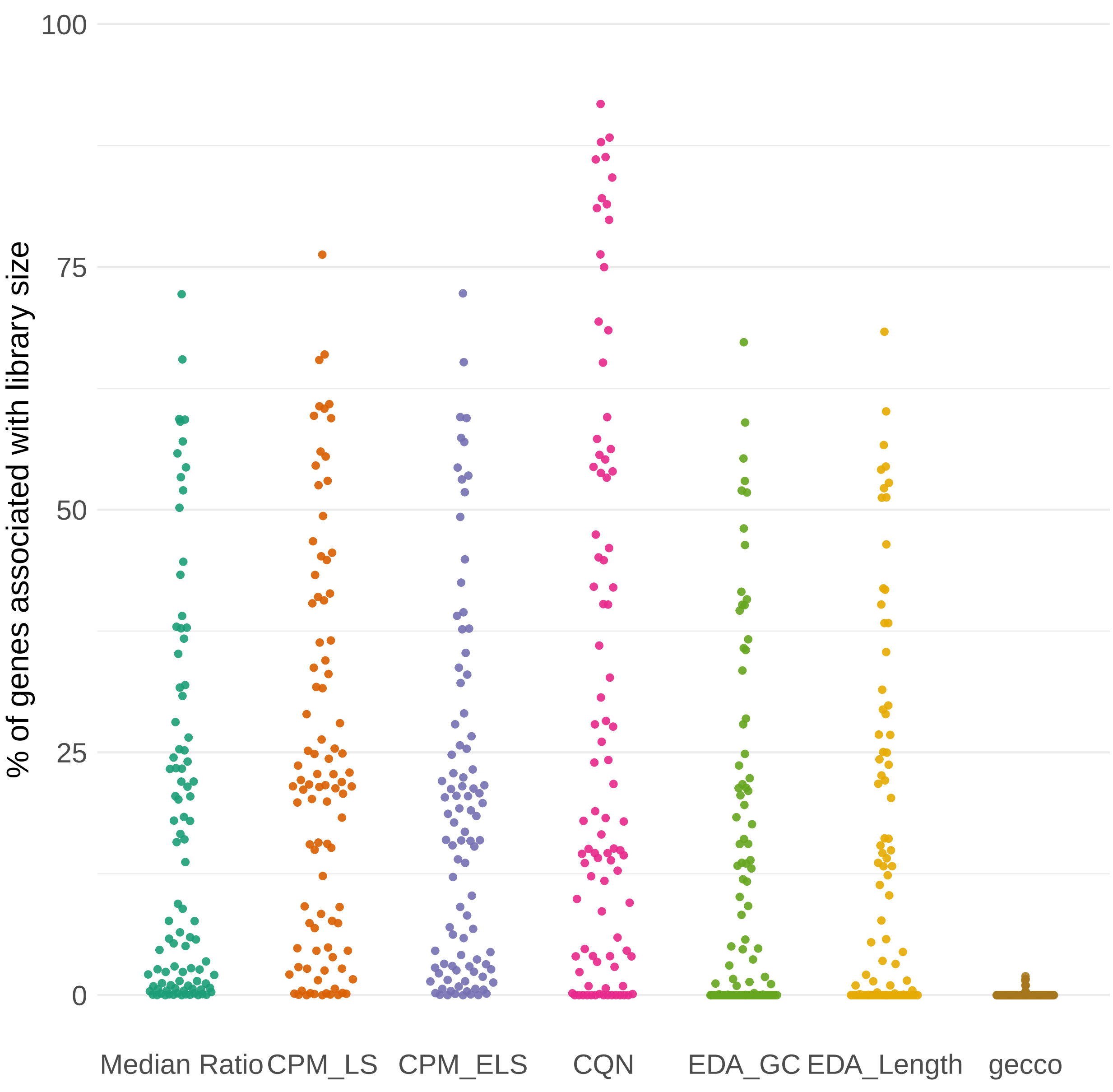


Supplementary Figure S4: Proportion of genes in each of the 102 analyzed datasets whose normalized counts remain correlated with library size under different normalization methods.

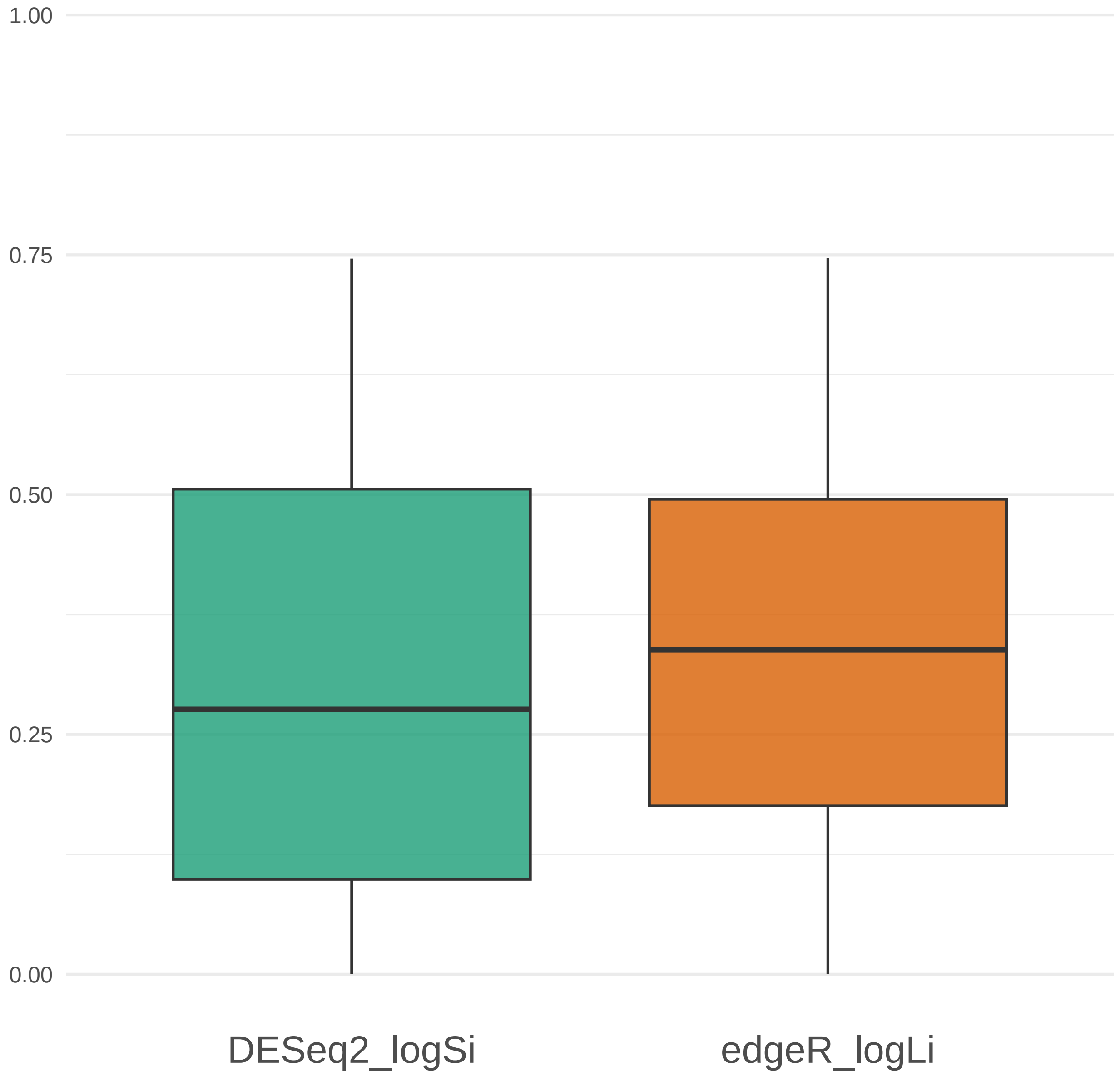


Supplementary Figure S5: Proportion of genes identified by differential expression (DE) analysis as significantly associated with sequencing-depth scaling factors across 102 public datasets. Panel (a) shows DESeq2 using $\log(s_i)$, and panel (b) shows edgeR using $\log(L_i)$. Significance is defined by BH-adjusted $p < 0.05$.

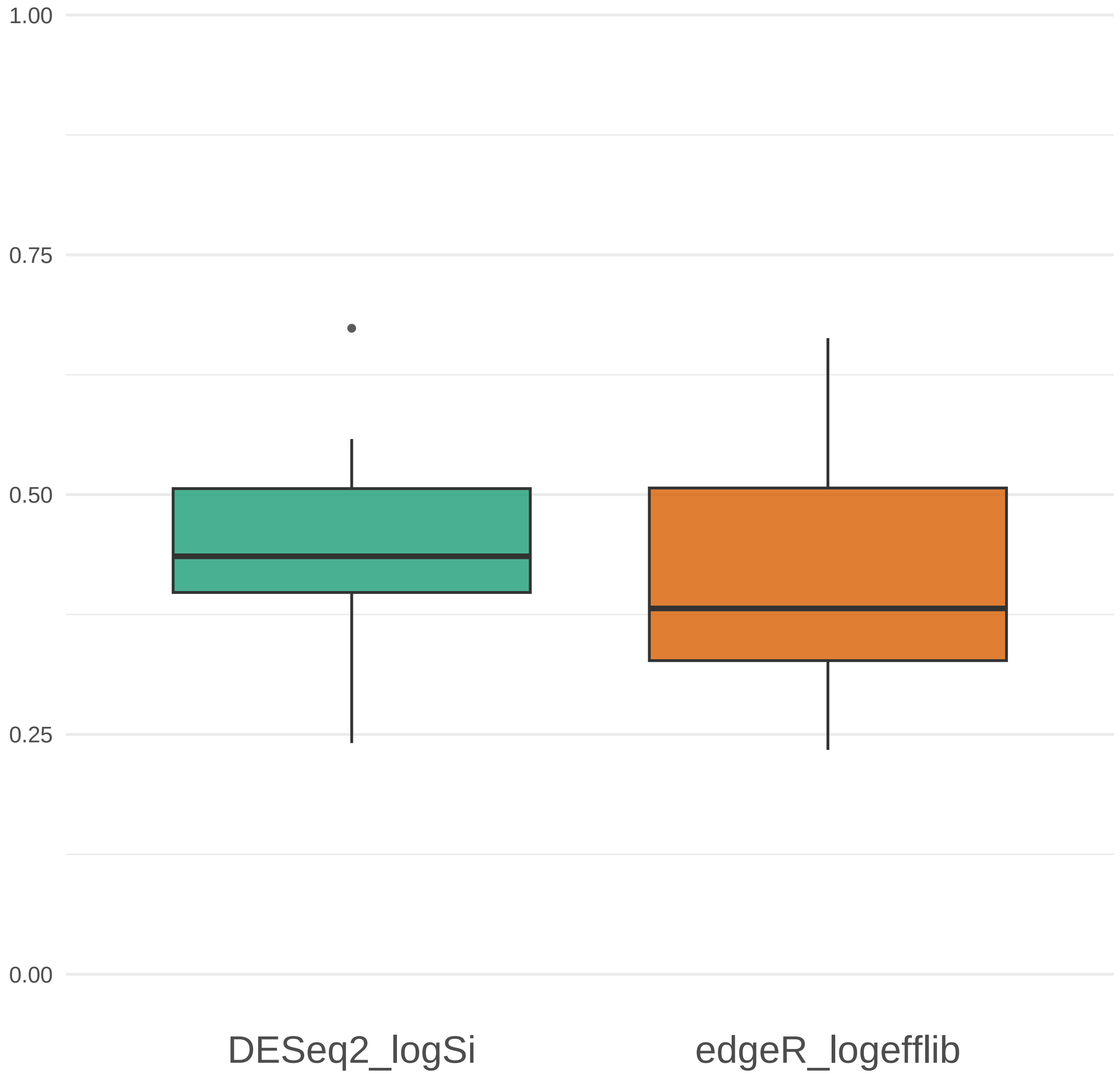


Supplementary Figure S6: Proportion of genes identified by differential expression (DE) analysis as significantly associated with sequencing-depth scaling factors after adjusting for clinical covariates across 8 in-house datasets. Panel (a) shows DESeq2 using $\log(s_i)$, and panel (b) shows edgeR using $\log(L_i)$. Significance is defined by BH-adjusted $p < 0.05$.

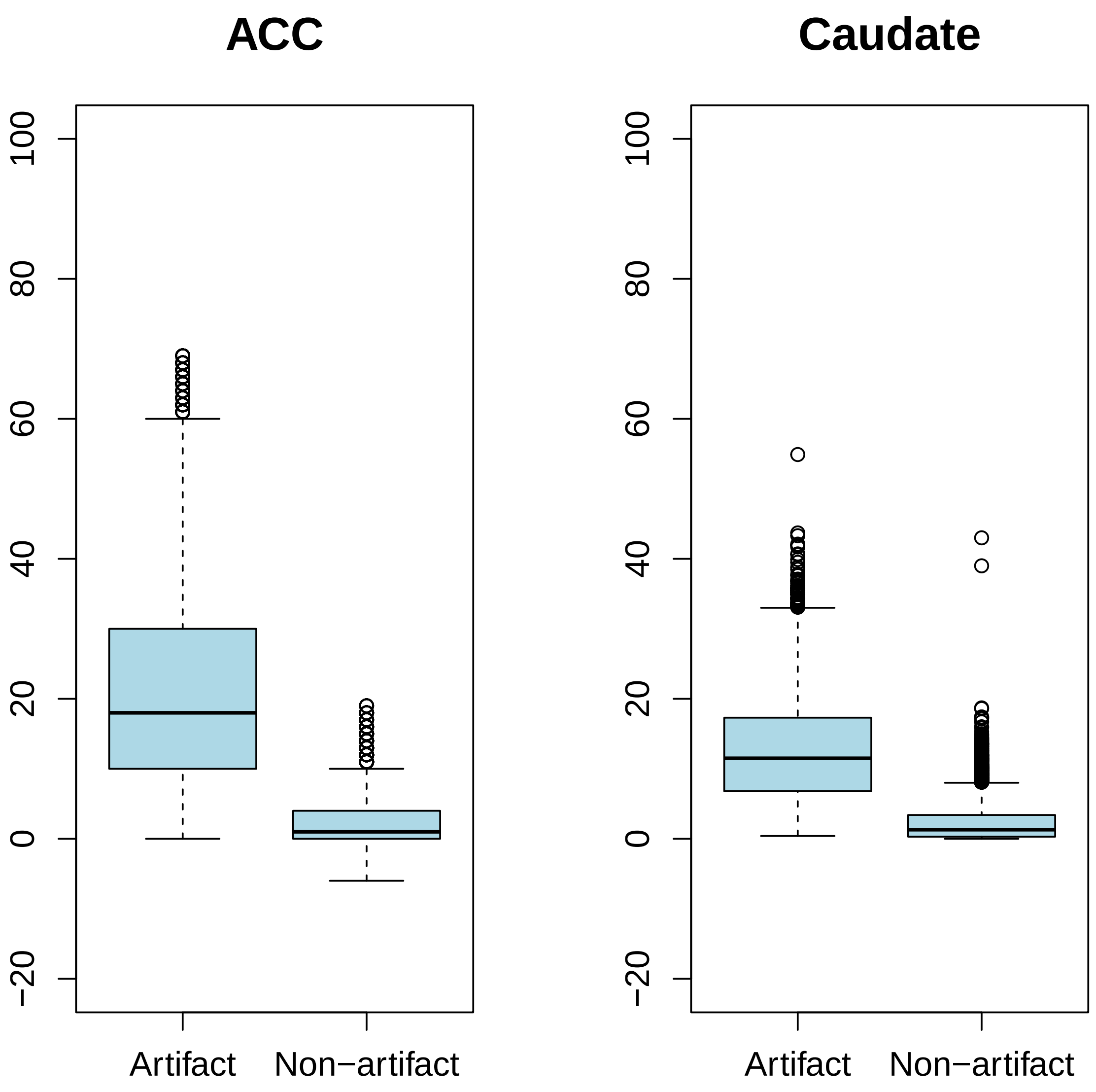


Supplementary Figure S7: ACC and Caudate RNA-seq Data: Differences in BIC Between *edgeR* and *gecco_edgeR* for artifact and Non-artifact Genes

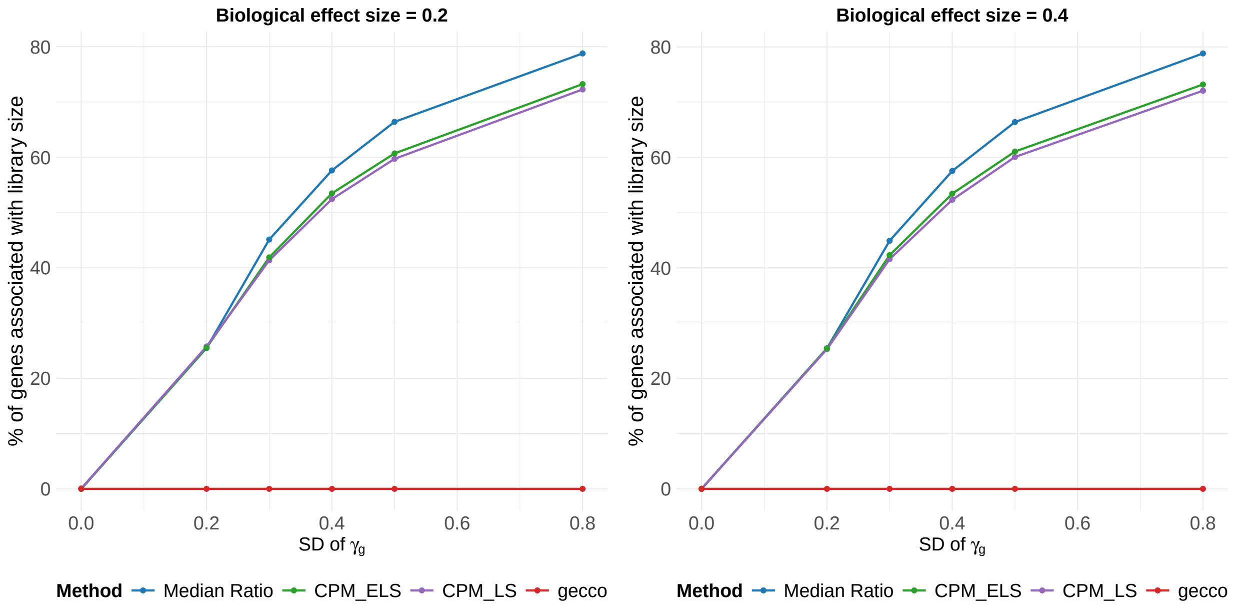


Supplementary Figure S8: Persistence of sequencing depth-associated correlations (BH-adjusted $p < 0.05$) across normalization methods and levels of $\gamma_g$, given a biological effect size of $\beta_{1,g}$. Panel (a): $\beta_{1,g} = \pm 0.2$. Panel (b): $\beta_{1,g} = \pm 0.4$.

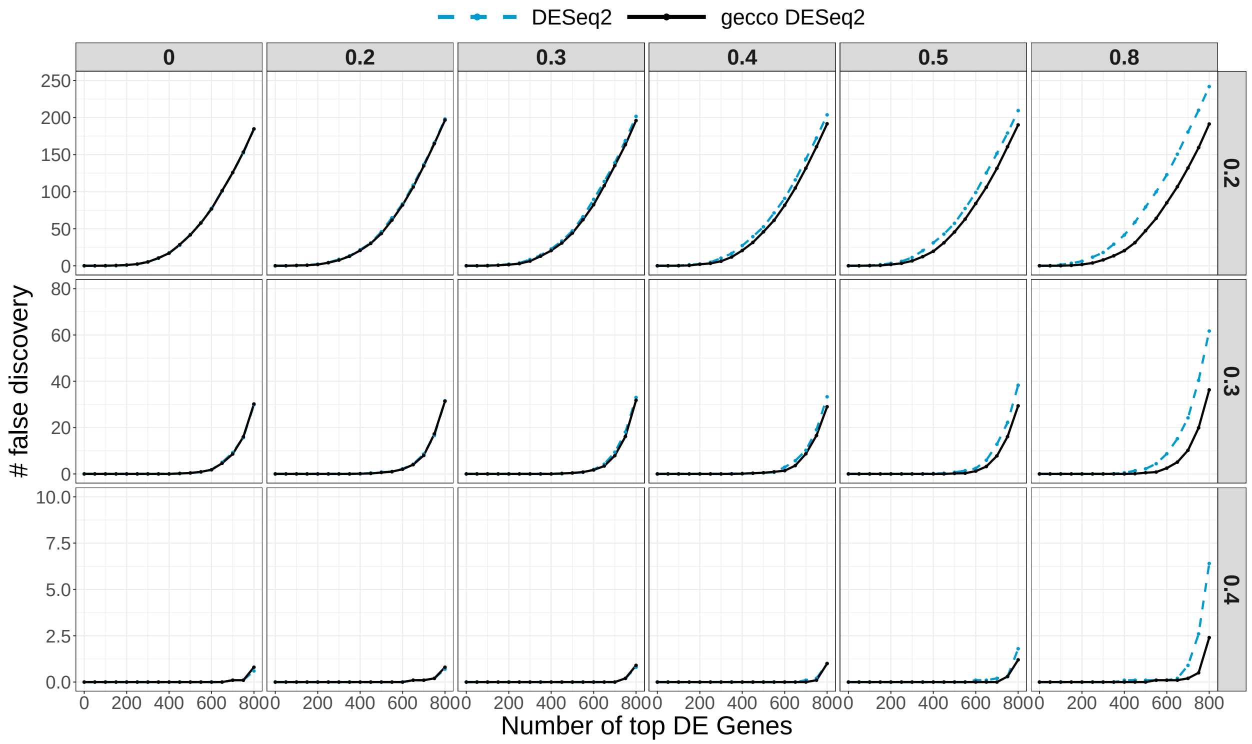


Supplementary Figure S9: False discovery counts for DESeq2 and gecco_DESeq2 across top-ranked differentially expressed genes under varying biological effect sizes and sequencing bias levels.

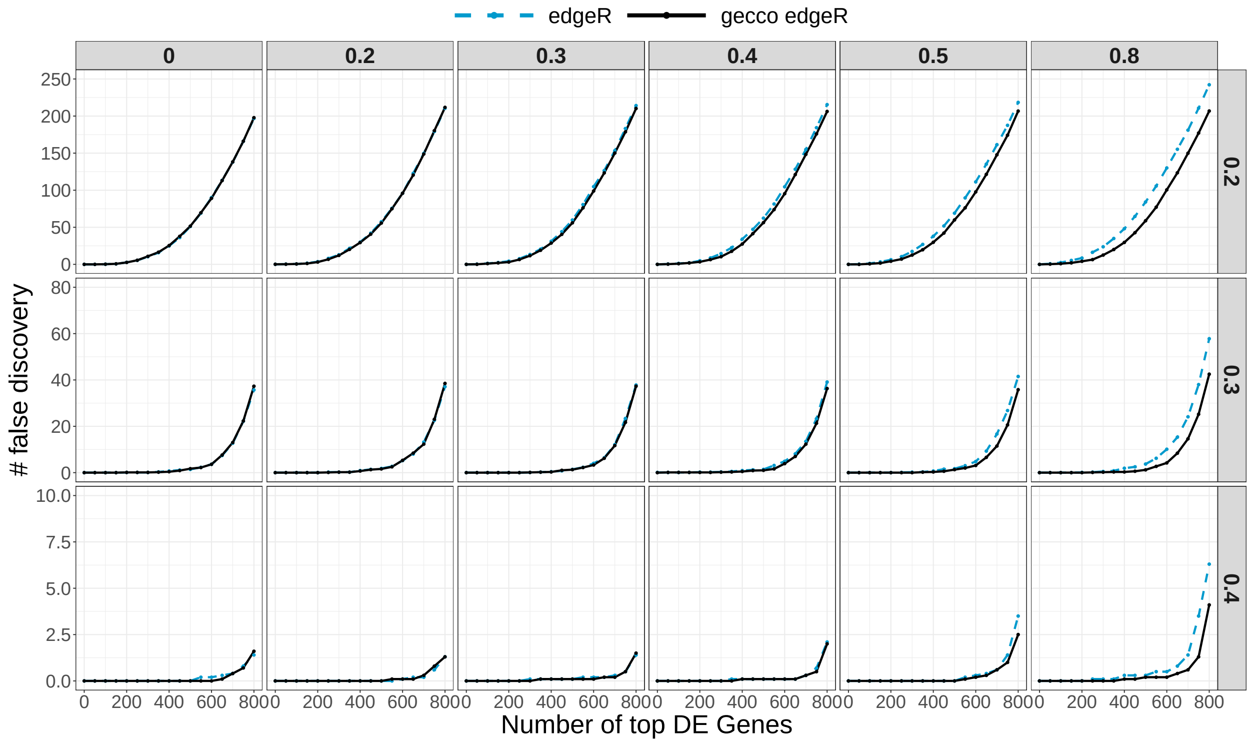


Supplementary Figure S10: False discovery counts for edgeR and gecco_edgeR across top-ranked differentially expressed genes under varying biological effect sizes and sequencing bias levels.

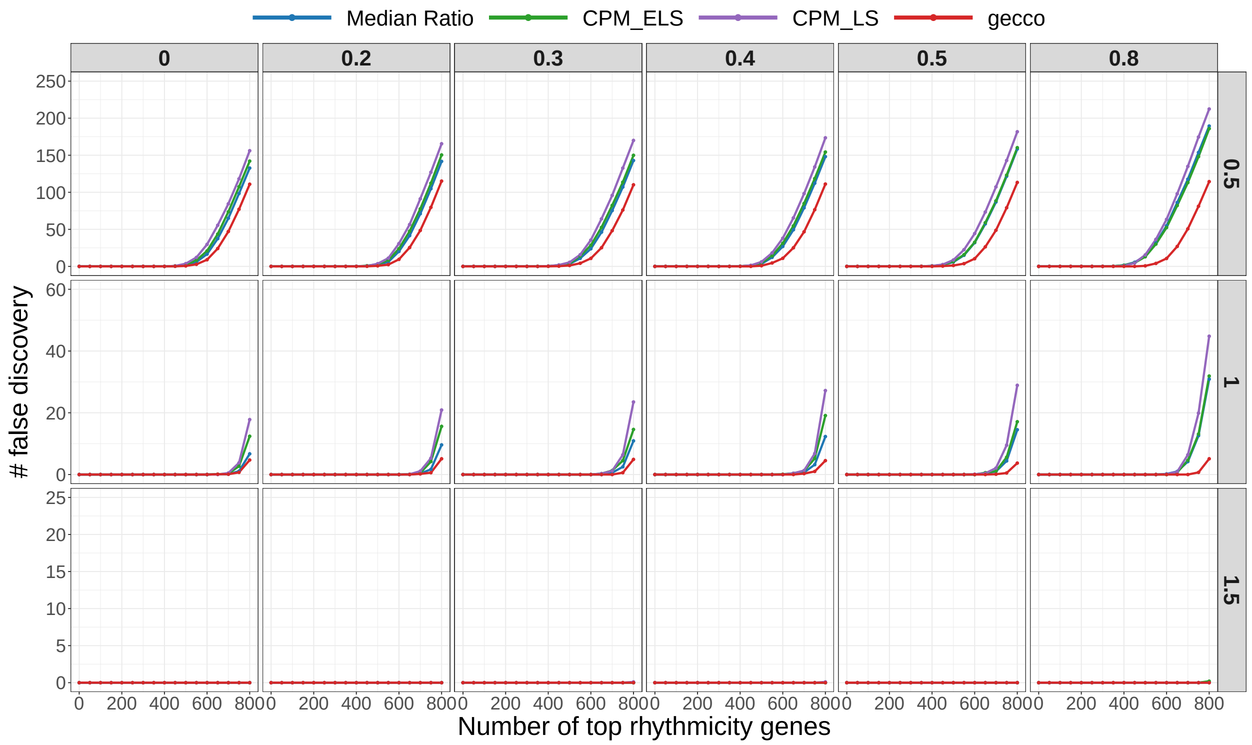


Supplementary Figure S11: False discovery counts across top-ranked circadian genes under four normalization methods.

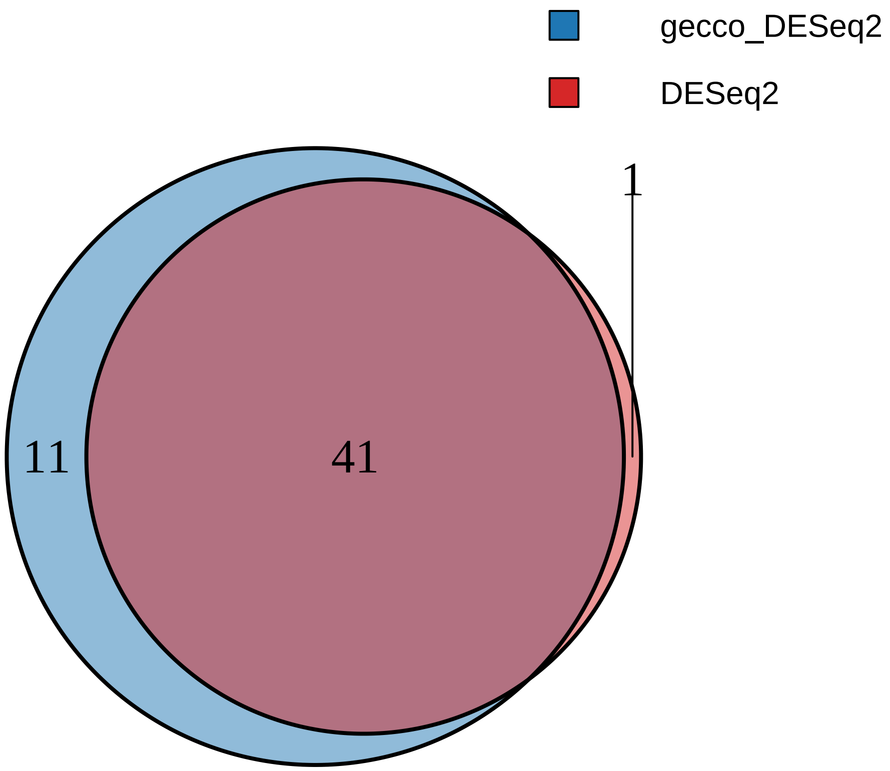


Supplementary Figure S12: Venn diagram of differentially expressed genes identified by DESeq2 and gecco_DESeq2.

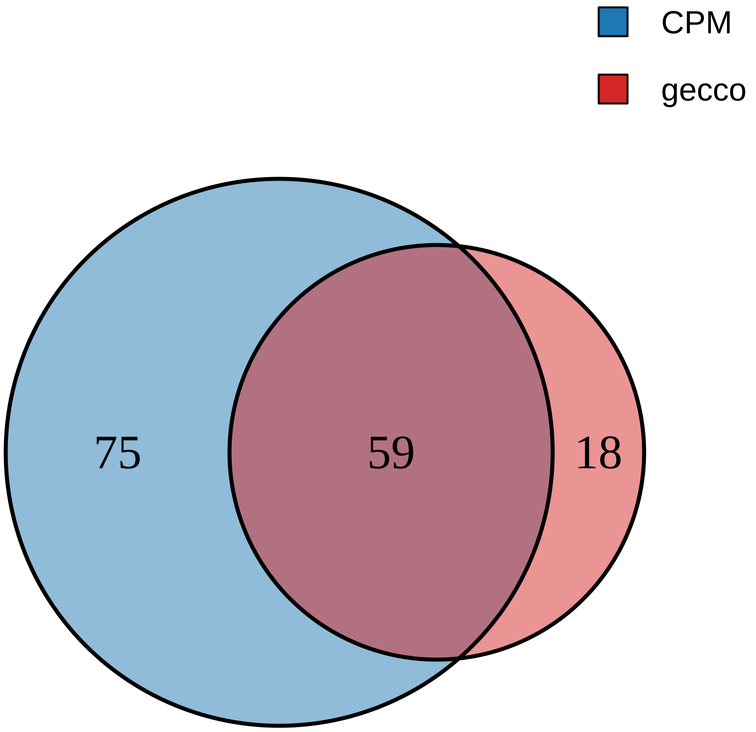


Supplementary Figure S13: Venn diagram of differentially expressed genes identified by CPM_LS and gecco.

#### 2 Supplementary Table1

Table S1: Summary of GEO and TCGA RNA-seq datasets used in this study.

| file | sample.size |
|---|---|
| GSE114407 | 195 |
| GSE114922 | 90 |
| GSE122380 | 297 |
| GSE123359 | 109 |
| GSE126409 | 104 |
| GSE131047 | 104 |
| GSE134874 | 242 |
| GSE136610 | 237 |
| GSE136825 | 103 |
| GSE139533 | 155 |
| GSE140244 | 200 |
| GSE142648 | 150 |
| GSE142840 | 89 |
| GSE143158 | 176 |
| GSE143897 | 122 |
| GSE145531 | 176 |
| GSE147352 | 118 |
| GSE149601 | 195 |
| GSE149809 | 100 |
| GSE153610 | 85 |
| GSE156063 | 234 |
| GSE156212 | 106 |
| GSE157585 | 135 |
| GSE163498 | 120 |
| GSE164649 | 151 |
| GSE167056 | 144 |
| GSE168017 | 94 |
| GSE168409 | 117 |
| GSE168698 | 122 |
| GSE169030 | 140 |
| GSE171343 | 240 |
| GSE172367 | 190 |
| GSE174482 | 134 |
| GSE174693 | 102 |
| GSE175384 | 111 |
| GSE179277 | 237 |
| GSE180707 | 100 |
| GSE181157 | 173 |
| GSE181859 | 221 |
| GSE182117 | 186 |
| GSE182622 | 270 |
| GSE189672 | 110 |
| GSE191084 | 192 |
| GSE195727 | 139 |
| GSE196126 | 172 |
| GSE197307 | 282 |
| GSE198000 | 96 |
| GSE199849 | 136 |
| GSE205430 | 100 |
| GSE81730 | 288 |
| GSE95446 | 191 |
| GSE111892 | 132 |

Table S1: Summary of GEO and TCGA RNA-seq datasets used in this study. *(continued)*

| file | sample.size |
| --- | --- |
| GSE117623 | 138 |
| GSE124284 | 286 |
| GSE125529 | 113 |
| GSE130633 | 126 |
| GSE141198 | 148 |
| GSE142174 | 108 |
| GSE144269 | 140 |
| GSE148025 | 198 |
| GSE154377 | 134 |
| GSE157194 | 166 |
| GSE160329 | 166 |
| GSE160521 | 177 |
| GSE161731 | 201 |
| GSE164416 | 133 |
| GSE164877 | 226 |
| GSE169031 | 263 |
| GSE174473 | 155 |
| GSE174659 | 138 |
| GSE184941 | 180 |
| GSE185512 | 180 |
| GSE186908 | 162 |
| GSE199452 | 111 |
| GSE201412 | 105 |
| GSE201533 | 161 |
| GSE201535 | 161 |
| GSE203395 | 294 |
| GSE206680 | 123 |
| GSE93326 | 204 |
| GSE95450 | 191 |
| TCGA_BLCA | 414 |
| TCGA_BRCA | 1119 |
| TCGA_COAD | 483 |
| TCGA_DLBC | 48 |
| TCGA_HNSC | 504 |
| TCGA_KICH | 66 |
| TCGA_KIRC | 542 |
| TCGA_KIRP | 291 |
| TCGA_LGG | 532 |
| TCGA_LUAD | 541 |
| TCGA_PRAD | 502 |
| TCGA_READ | 167 |
| TCGA_SKCM | 472 |
| TCGA_STAD | 420 |
| TCGA_THCA | 513 |
| TCGA_UCEC | 554 |
| TCGA_ACC | 79 |
| TCGA_CESC | 306 |
| TCGA_GBM | 170 |
| TCGA_LAML | 178 |
| TCGA_LIHC | 374 |
| TCGA_LUSC | 502 |
| TCGA_OV | 430 |
| TCGA_UCS | 57 |

### 3 Supplementary Table2

Table S2: Summary of datasets used for differential expression analyses.

| Dataset | Type | Sample size | Description |
|---|---|---|---|
| in-house1 | in-house | 239 | Human anterior cingulate cortex (ACC) samples from control, major depressive disorder (MDD), Schizophrenia (SCZ) and Biopolar (BP), including layer 3 and layer 5 brain regions. |
| in-house2 | in-house | 79 | Human anterior cingulate cortex (ACC) samples from control subjects, including layer 3 and layer 5 brain regions. |
| in-house3 | in-house | 86 | Human nucleus accumbens (NAc) samples from major depressive disorder (MDD) and control subjects. |
| in-house4 | in-house | 248 | Human anterior cingulate cortex (ACC) samples from control, major depressive disorder (MDD), Schizophrenia (SCZ) and Biopolar (BP) subjects. |
| in-house5 | in-house | 249 | Human dorsolateral prefrontal cortex (DLPFC) samples from control, major depressive disorder (MDD), Schizophrenia (SCZ) and Biopolar (BP) subjects. |
| GSE160521 | GEO public | 59 | Human Caudate brain region samples from control subjects. |
| GSE160521 | GEO public | 59 | Human nucleus accumbens (NAc) brain region samples from control subjects. |
| GSE160521 | GEO public | 59 | Human Putamen brain region samples from control subjects. |
